# Integrated CHARGE syndrome models reveal epigenetic modulators of reproductive phenotypes

**DOI:** 10.64898/2026.03.31.715488

**Authors:** Federica Amoruso, Federica La Rocca, Pamela Santonicola, Alyssa J.J. Paganoni, Giuseppina Zampi, Stefano Manzini, Fabrizio Fontana, Riccardo Cristofani, Roberto Oleari, Elia Di Schiavi, Anna M. Cariboni

## Abstract

Loss-of-function variants in *CHD7* cause CHARGE syndrome (CS), a rare developmental disorder showing multisystem malformations, including reproductive defects linked to gonadotropin-releasing hormone (GnRH) neuron dysfunction. CHD7 encodes a chromatin remodeler essential for early transcriptional regulation across various tissues. Currently, no pharmacological treatments exist, and approaches aimed at identifying tissue-specific CHD7 targets are also lacking, making CS treatment an unmet clinical need.

To explore mechanisms relevant to CS-associated reproductive defects, we established a dual screening platform combining CRISPR-engineered *Chd7*-depleted mouse GnRH neurons with a *Caenorhabditis elegans chd-7* mutant showing reproductive abnormalities. Transcriptomic and functional analyses of *Chd7*-deficient cells revealed impaired cellular processes along with dysregulation of *semaphorin* (*Sema*) genes, key regulators of GnRH neuron development. A screen of 234 epigenetic modulators in *C. elegans* identified compounds modifying aberrant mutant phenotype, two of which also rescued cellular defects and *Sema* expression in vitro.

Altogether, these findings indicate that CHD7 deficiency reshapes neuroendocrine-relevant pathways and that selected compounds modulate CS-associated phenotypes across species, with SEMA signalling emerging as candidate druggable downstream pathway in CS requiring further mechanistic validation.

## Introduction

CHARGE syndrome (CS) is a rare (∼1:15,000 births) genetic disease characterized by a broad spectrum of congenital malformations with variable severity and expressivity. Clinical features include congenital heart defects, choanal atresia, growth retardation, genital malformations, developmental delay, eye coloboma, ear anomalies, and deafness (Pagon *et al*, 1981; Issekutz *et al*, 2005). Neurological manifestations such as intellectual disability, epilepsy, autism spectrum traits, and neuroendocrine defects, including hypogonadotropic hypogonadism (HH), are also frequent (Pinto *et al*, 2005; Balasubramanian & Crowley, 2017; Thomas *et al*, 2022; Abadie *et al*, 2020). CS is associated with high morbidity and mortality, with approximately 30% of affected individuals dying before the age of five, often following multiple surgical interventions to correct life-threatening anomalies (Blake *et al*, 1990, 1998; Trider *et al*, 2017). Despite advances in clinical management, no pharmacological treatments are currently available, underscoring a critical unmet therapeutic need (van Ravenswaaij-Arts & Martin, 2017; Hsu *et al*, 2014).

Heterozygous loss-of-function variants in chromodomain helicase DNA binding protein 7 (*CHD7*) gene account for more than 85% of CS cases (Ufartes *et al*, 2021). *CHD7* encodes an ATP-dependent chromatin remodeler that plays a pivotal role in regulating gene expression during embryonic development (Basson & van Ravenswaaij-Arts, 2015; Lettieri *et al*, 2021). Genome-wide analyses have shown that CHD7 is predominantly recruited to active enhancers and promoters, where it facilitates chromatin opening and transcriptional activation. In a context-dependent manner, CHD7 can also promote chromatin compaction and transcriptional repression, depending on developmental stage and tissue-specific cues (Schnetz *et al*, 2010, 2009; Zentner *et al*, 2011; Feng *et al*, 2017b). Most *CHD7* variants identified in CS patients arise *de novo* and impair nucleosome remodelling, leading to altered chromatin accessibility for transcription factors (Jongmans *et al*, 2006; Bouazoune & Kingston, 2012). Notably, pathogenic *CHD7* variants have also been reported in Kallmann syndrome, a developmental disorder characterized by HH and anosmia, highlighting shared molecular mechanisms underlying reproductive phenotypes across related conditions (Topaloğlu, 2017; Dodé & Hardelin, 2010; Gonçalves *et al*, 2019).

CHD7 is highly expressed in the developing nervous system, particularly within the neuroectoderm and neural crest derivatives, including cranial nerves and facial mesenchyme (Bosman *et al*, 2005; Sanlaville *et al*, 2006). CHD7 is required for neural progenitor patterning, neural stem cell maintenance, gonadotropin-releasing hormone (GnRH) neuron specification, and neural crest cell migration and differentiation (Bajpai *et al*, 2010; Chai *et al*, 2018; Jones *et al*, 2015). Disruption of these processes results in major neurodevelopmental abnormalities, including cerebellar hypoplasia, microcephaly, ventriculomegaly, midline defects, aberrant GnRH neuron neurogenesis and migration, as well as olfactory bulb and corpus callosum malformations, all hallmark features of CS (Feng *et al*, 2017a; Huang *et al*, 2025; Balasubramanian & Crowley, 2017; Pinto *et al*, 2005; Layman *et al*, 2011).

While gene replacement strategies could, in principle, represent a curative approach for CS, their clinical translation is hampered by major technical challenges, including the large size of the *CHD7* gene, the need for broad tissue tropism, and the requirement for early, potentially prenatal, delivery without maternal risk (van Ravenswaaij-Arts & Martin, 2017). Consequently, recent efforts have explored pharmacological modulation of CHD7 downstream pathways as a more feasible postnatal therapeutic strategy. These studies have provided important proof-of-concept that CS-associated phenotypes can be pharmacologically modified, including through retinoic acid signalling modulation in *Chd7*-mutant mice (Micucci *et al*, 2013), phenotype-based compound screening in *chd7*-deficient zebrafish and nematodes (Jamadagni *et al*, 2021), and epigenetic rebalancing via EZH2 inhibition in human brain organoids and mice (Huang *et al*, 2025).

In this context, the *Caenorhabditis elegans* (*C. elegans*) *chd-7*(*gk290)* allele represents the most widely used in vivo model of *CHD7* loss of function (Schmeisser *et al*, 2017; Jamadagni *et al*, 2021; Rawsthorne *et al*, 2021; Jofré *et al*, 2022). The gk290 deletion removes the BRK domain and the C-terminal region of the protein, generating a null allele that mirrors the functional relevance of this domain in humans (Hale *et al*, 2016). Mutant animals display impaired locomotion, reduced lifespan, and GABAergic defects (Jamadagni *et al*, 2021), and additional autism-associated missense variants, including the CS mutation L1257R, introduced into the *chd-7* locus further disrupt fertility and development (Wong *et al*, 2019). These features make *C. elegans* a genetically tractable and phenotypically robust system for dissecting CHD7-dependent pathways and for phenotype-based compound screening.

Despite the strong genetic association between *CHD7* haploinsufficiency and CS, affected individuals display marked phenotypic variability both in the number and severity of clinical manifestations (Jongmans *et al*, 2008), suggesting that genetic modifiers and downstream signalling pathways critically shape disease expression in a *CHD7*-deficient background (Basson & van Ravenswaaij-Arts, 2015).

Among candidate modifier pathways, semaphorins (SEMA) – a family of axon guidance molecules with key roles in neural and reproductive development, and their receptors plexins and neuropilins (PLXN/NRP) – have emerged as relevant CHD7-regulated targets (Lettieri *et al*, 2021). CHD7 directly or indirectly regulates the expression of multiple *SEMA* genes, including *Sema3a* (Schulz *et al*, 2014) and *Sema6c* (Qu *et al*, 2002), which are widely expressed in the developing brain and exert crucial neurodevelopmental functions (Oleari *et al*, 2019). Moreover, pathogenic variants in class 3 semaphorins *SEMA3A* and *SEMA3E* have been identified in patients with CS and/or neuroendocrine forms of reproductive failure (Lalani *et al*, 2004; Cariboni *et al*, 2015; Ufartes *et al*, 2018). Genetic interaction studies further support a functional interplay between *CHD7* and *SEMA* signalling in pathways essential for GnRH neuron development (Cariboni *et al*, 2015).

Despite accumulating evidence linking CHD7 dysfunction to SEMA signalling, the therapeutic potential of modulating this pathway in CS remains unexplored. Here, we adopted epigenetic modulation as a discovery-driven strategy to interrogate downstream pathways associated with *CHD7* deficiency, with particular attention to SEMA signalling as a genetically supported candidate. Our aim was to identify pathway-level mechanisms that may contribute to GnRH neuron development and neuroendocrine reproductive dysfunction in CS.

To explore whether pharmacological intervention could mitigate phenotypes associated with CHD7 loss, we combined complementary in vitro and in vivo models of CS in a cross-species screening framework. Rather than directly targeting a predefined effector, our approach sought to identify small molecules capable of modulating disease-relevant cellular and reproductive phenotypes, and to subsequently assess their impact on candidate downstream pathways, including SEMA signalling.

Using CRISPR/Cas9, we generated *Chd7*-depleted cell clones in the murine immortalized GnRH neuron cell line GN11, which revealed defects in cell migration, adhesion and proliferation associated with altered *SEMA* gene expression. In parallel, we employed the *Caenorhabditis elegans* (*C. elegans*) *chd-7*(*gk290)* mutant, characterized by a readily quantifiable reproductive defect, to establish a semi-automated, phenotype-based screening platform. Screening of an epigenetic compound library identified five candidates that ameliorated the reproductive phenotype in mutant worms. Selected compounds also improved cellular defects in *Chd7*-depleted GN11 cells and were associated with partial restoration of *Sema* gene expression.

Collectively, these findings establish a cross-species framework for identifying small molecules that compensate for *CHD7* loss by modulating downstream SEMA pathways, with potential translational relevance for CS and related reproductive disorders.

## Results

### CRISPR/Cas9 genome editing establishes *Chd7*-deficient GN11 cells for in vitro CS modelling

To investigate *Chd7-Sema* interactions in vitro, we employed CRISPR/Cas9-mediated genome editing in GN11 cells, a murine model immature migrating GnRH neurons widely used to study neurodevelopmental processes (Maggi *et al*, 2000; Cariboni *et al*, 2011a, 2015, 2007). GN11 cells express *Chd7* together with multiple SEMA genes, including *Sema3a*, *Sema3c*, *Sema3f*, and their receptors (*Neuropilin1-2* and *PlexinA1-4*), making them a suitable system to interrogate *Chd7*-dependent regulation of SEMA signalling in GnRH neurons (Kim *et al*, 2008; Cariboni *et al*, 2007). To generate a *Chd7*-depleted model, we designed two single guide RNAs (sgRNAs) targeting exon 2 and exon 38 of the murine *Chd7* gene, enabling the excision of the intervening coding region, and causing the consequent loss of protein expression (**Fig. 1A**). sgRNAs were cloned into a plasmid encoding the *Streptococcus pyogenes* Cas9 (SpCas9) nuclease and co-transfected into GN11 cells. PCR screening on genomic DNA (gDNA) with custom target exon-flanking and internal oligonucleotides identified six independent clones with a heterozygous *Chd7* deletion, hereafter referred to as *Chd7*-knockout (*Chd7*-KO) clones. These clones showed evidence of gene deletion on one allele, as indicated by the presence of a deletion-specific band in the gDNA PCR, while retaining exon 3 from the non-targeted allele, thus confirming that CRISPR/Cas9-mediated editing occurred on a single allele only (**Fig. EV1A,B**). Gene editing was further confirmed by Sanger sequencing (**Fig. EV1C**). The monoallelic deletion obtained models the haploinsufficiency typically observed in CS, making these clones a reliable in vitro disease model.

**Figure 1.**
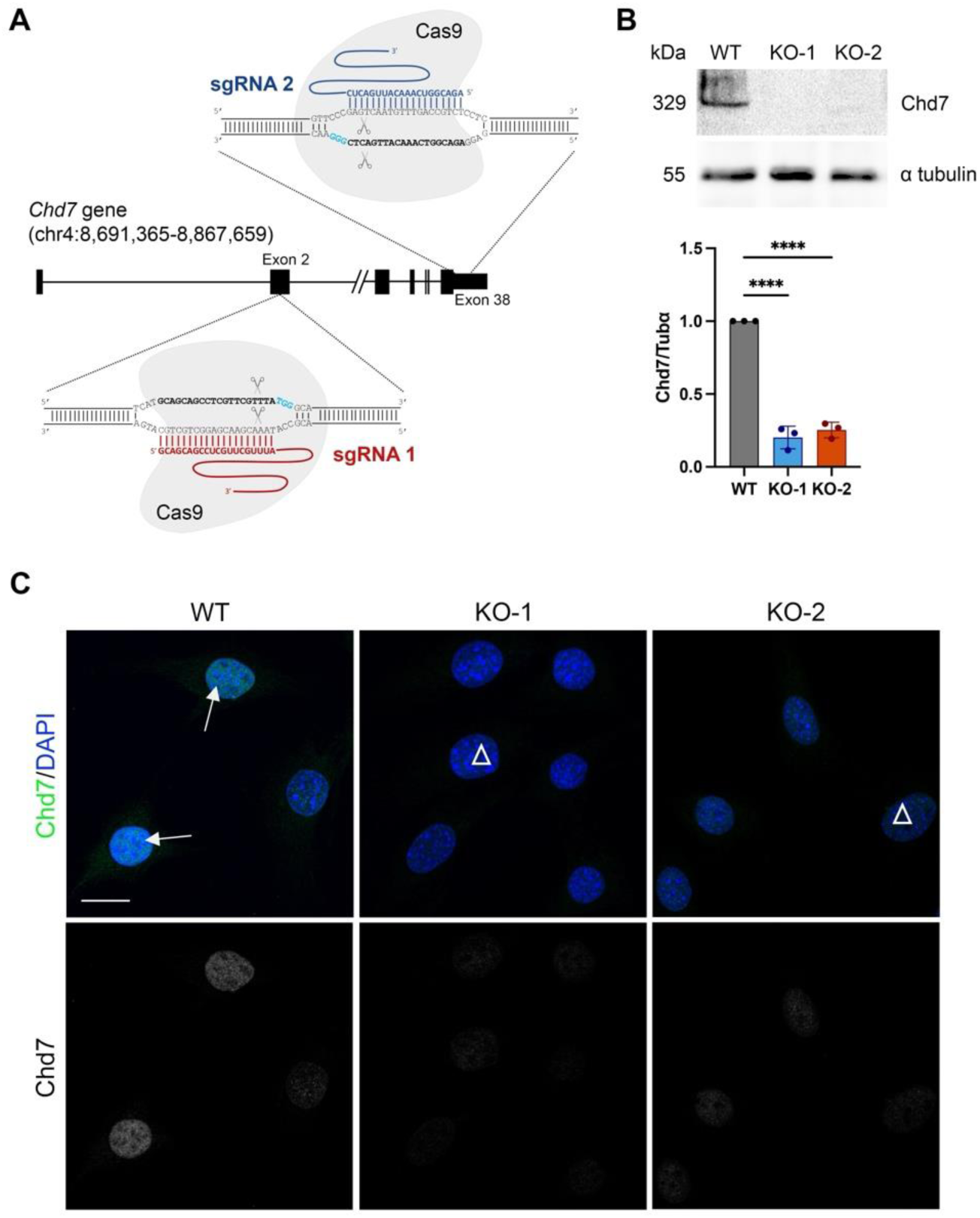
Generation of *Chd7*-KO cell lines using CRISPR/Cas9 genome editing technique. (A) Schematic representation of CRISPR/Cas9 genome editing strategy targeting the murine *Chd7* gene. Single guide RNA (sgRNA) sequences specific to exon 2 (red) and exon 38 (blue) are depicted. (B) Western Blot analysis of Chd7 protein expression in WT and CRISPR-edited GN11 cell clones (KO-1 and KO-2). α tubulin was used as loading control. Data are presented as mean ± SD (N = 3; One-way ANOVA followed by Dunnett’s multiple comparisons test, **** p < 0.0001). (C) Representative confocal microscopy images of GN11 WT and *Chd7*-KO cells following immunocytochemistry with anti-Chd7 antibody (green above, grey below). Nuclei were counterstained with DAPI (blue). Scale bar: 25 μm. Abbreviations: chr, chromosome; SANT, SWI3, ADA2, N-CoR, and TFIIIB; BRK, Braham and Kismet;M, marker; NTC, non template control; Tubα, tubulin alpha.

To confirm loss of protein expression, Chd7 levels were assessed by Western blot and immunocytochemistry, revealing variable depletion across clones. Two independent clones, with significantly reduced or undetectable Chd7 protein levels compared to wild-type (WT) were selected for downstream experiments and are hereafter referred to as KO-1 and KO-2 (**Fig. 1B,C**).

### Transcriptomic profiling reveals dysregulation of cell cycle, migration, and semaphorin signalling pathways upon *Chd7* depletion in GN11 cells

To investigate the transcriptomic consequences of *Chd7* depletion, we performed bulk RNA sequencing (RNA-seq) on KO-1, KO-2 and WT GN11 cells (three biological replicates per experimental group). Principal component analysis (PCA) showed strong clustering of replicates within each group, with KO samples clearly separating from WT, indicating consistent transcriptional profiles within genotypes (**Fig. 2A**). In this analysis, 1748 and 1554 differentially expressed genes (DEGs; |log₂FC| > 1, adjusted *p-value* < 0.05) were identified in the WT vs KO-1 and WT vs KO-2 comparisons, respectively. Among them, 704 DEGs (435 upregulated and 269 downregulated genes) were shared between KO-1 and KO-2, with most of them having the same expression trend (i.e. up- or down-regulated in both conditions) (**Fig. 2B,C**). Notably, the analysis of DEGs shared by both *Chd7*-KO clones identified several genes previously implicated in GnRH neuron development through direct experimental evidence, including *Slit2*, *Camk2a* and *Wls* (Cariboni *et al*, 2012; Lund *et al*, 2016; Melamed *et al*, 2012; Wang *et al*, 2022), and genes known as pathogenic in patients with HH or CS, such as *Slit2*, *Alx4*, *Sema3f*, *Dlx5*, *Sema3a* and *Plxnb1* (Wu *et al*, 2022; Kayserili *et al*, 2009; Kotan *et al*, 2021; Joo *et al*, 2025; Zhou *et al*, 2018; Dai *et al*, 2020; Poch *et al*, 2024; Känsäkoski *et al*, 2014; Hanchate *et al*, 2012; Ufartes *et al*, 2018; Welch *et al*, 2022). Beyond neuroendocrine-relevant genes, shared DEGs also included regulators of epigenetic processes, such as *Sap30* and *Zfp979* (Zhang *et al*, 1998; Bertozzi *et al*, 2020), as well as genes involved in cytoskeletal organization, cell growth, motility and focal adhesions dynamics, including *Arhgap11a*, *Negr1*, *Mical1*, *Srgap3*, *Itga8* and *Iglon5* (Quadri *et al*, 2023; Szczurkowska *et al*, 2018; Terman *et al*, 2002; Dazzo *et al*, 2018; Bacon *et al*, 2013; Supriyanto *et al*, 2013; Salluzzo *et al*, 2023; Karis *et al*, 2018) (**Fig. 2D**).

**Figure 2.**
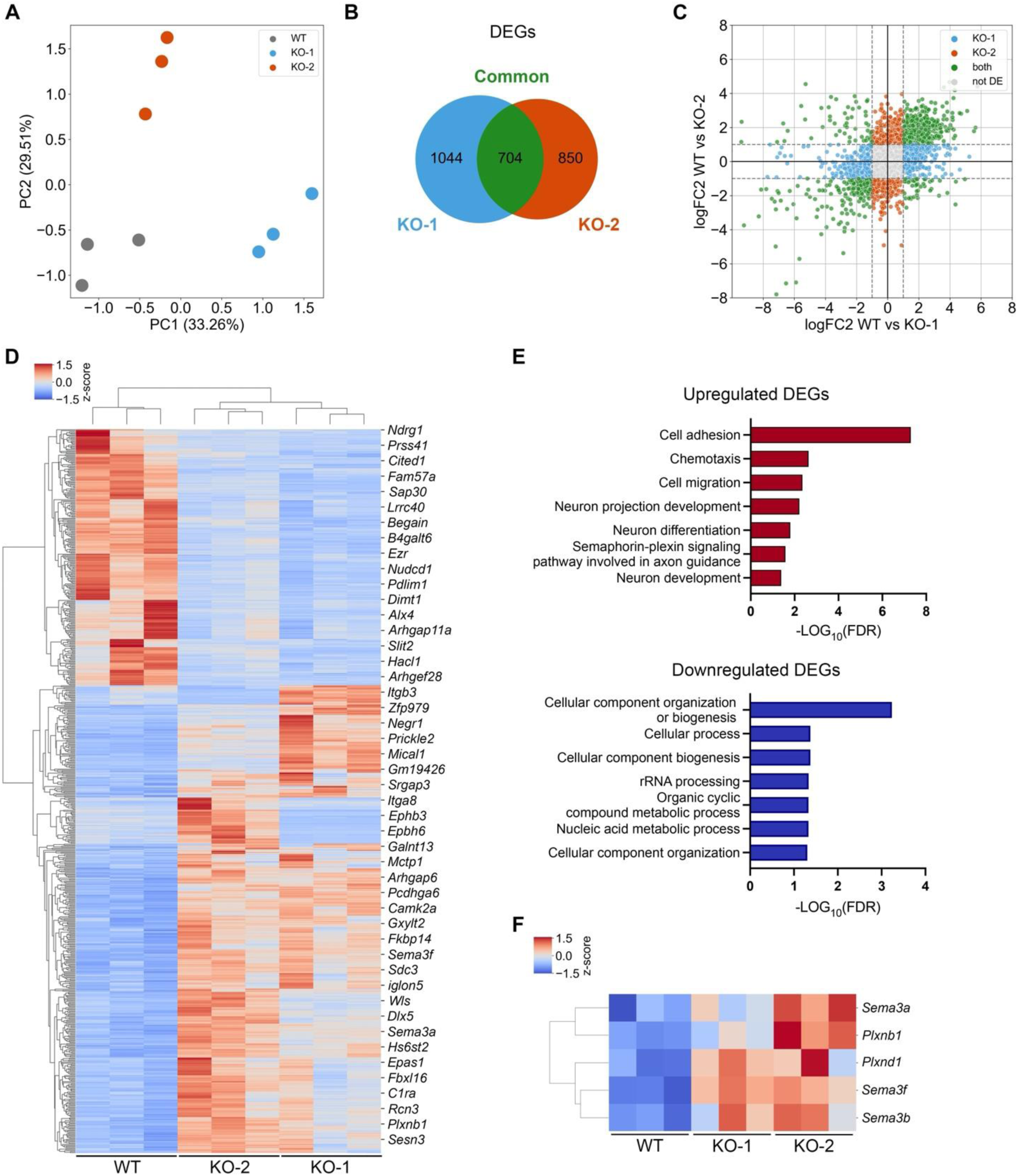
Transcriptomic profiling of WT and *Chd7*-KO GN11 cells. (A) Principal component analysis (PCA) of bulk RNA-seq data from WT and *Chd7*-KO (KO-1 and KO-2) GN11 cells, performed on annotated genes. The first two principal components (PC1 and PC2) are shown. (B) VENN diagram showing the overlap of differentially expressed genes (DEGs; |log_2_FC| > 1 and adj. *p-value* < 0.05) identified in WT vs KO-1 and WT vs KO-2 comparisons. (C) Volcano plot displaying differential gene expression in WT vs KO-1 and WT vs KO-2 comparisons. Significantly upregulated and downregulated genes (|log_2_FC| > 1 and adj. *p-value* < 0.05) are indicated. (D) Hierarchical clustering of z-scored gene expression values for DEGs (|log_2_FC| > 1 and adjusted *p-value* < 0.05) common to both WT vs KO-1 and WT vs KO-2 comparisons, using Euclidean distance metric (N = 3 biological replicates per experimental group). (E) Gene Ontology (GO) Biological Processes enrichment analysis performed using STRING on up- (top) and down-regulated (bottom) DEGs (FDR < 0.05). Enrichment scores are reported as -Log_10_FDR. (F) Heatmap of z-scored expression values for *Sema* and *Plxn* family genes differentially expressed between WT vs *Chd7*-KO GN11 cells (|Log_2_FC| > 1, adjusted *p-value* < 0.05). Abbreviations: PC, principal component; DEGs, differentially expressed genes; FDR, false discovery rate.

To get insights on the biological relevance of these transcriptional changes, we performed a functional enrichment analysis using reString (Manzini *et al*, 2021) based on Gene Ontology (GO) Biological Process annotations. Upregulated genes were significantly enriched in pathways related to cell adhesion, cell migration and chemotaxis (**Fig. 2E**, top). Although these pathways are associated with increased motility, a closer inspection revealed that many of the upregulated genes act as negative regulators of migration (e.g., *Sema3a*). Strikingly, one of the most significantly enriched pathways was the Semaphorin-Plexin signalling pathway involved in axon guidance, supporting a regulatory role for *Chd7* in the regulation of SEMA signalling. Conversely, downregulated genes were significantly enriched in pathways related to the cell cycle, cellular component biogenesis, and DNA/protein synthesis (**Fig. 2E**, bottom), consistent with a potential role of *Chd7* in supporting GN11 proliferation.

Notably, we identified five *Sema/Plxn* genes consistently upregulated in both *Chd7*-KO clones compared to WT GN11 cells, reinforcing the idea that CHD7 modulates SEMA signalling at the transcriptional level (**Fig. 2F**). This upregulation was confirmed by qPCR (**Fig. EV2A**).

### *Chd7* depletion impairs cell proliferation, migration and adhesion in GN11 cells

To investigate whether the transcriptomic alterations observed upon *Chd7* depletion in GN11 cells translate into functional phenotypic changes, we performed an array of in vitro assays focusing on cell proliferation and migration.

CHD7 has previously been implicated in the regulation of proliferation in neural stem cells and neurons (Jones *et al*, 2015; Layman *et al*, 2009; Feng *et al*, 2013; Ohta *et al*, 2016; Yao *et al*, 2020), suggesting a potentially conserved role in maintaining proliferative capacity in neurodevelopmental contexts. Thus, we monitored cell metabolic activity at 24, 48 and 72 hours (h) after seeding, using MTT assays. Both KO-1 and KO-2 clones exhibited a significant and progressive reduction in metabolic activity compared to WT GN11 cells at all time points considered (**Fig. 3A** and **Fig. EV2B**). Since a significant difference was still evident even 72 h after cell seeding, we chose this time point as the reference for proliferation assays, also with the perspective of assessing a potential rescue effect following extended drug treatment (**Fig. 3A**).

**Figure 3.**
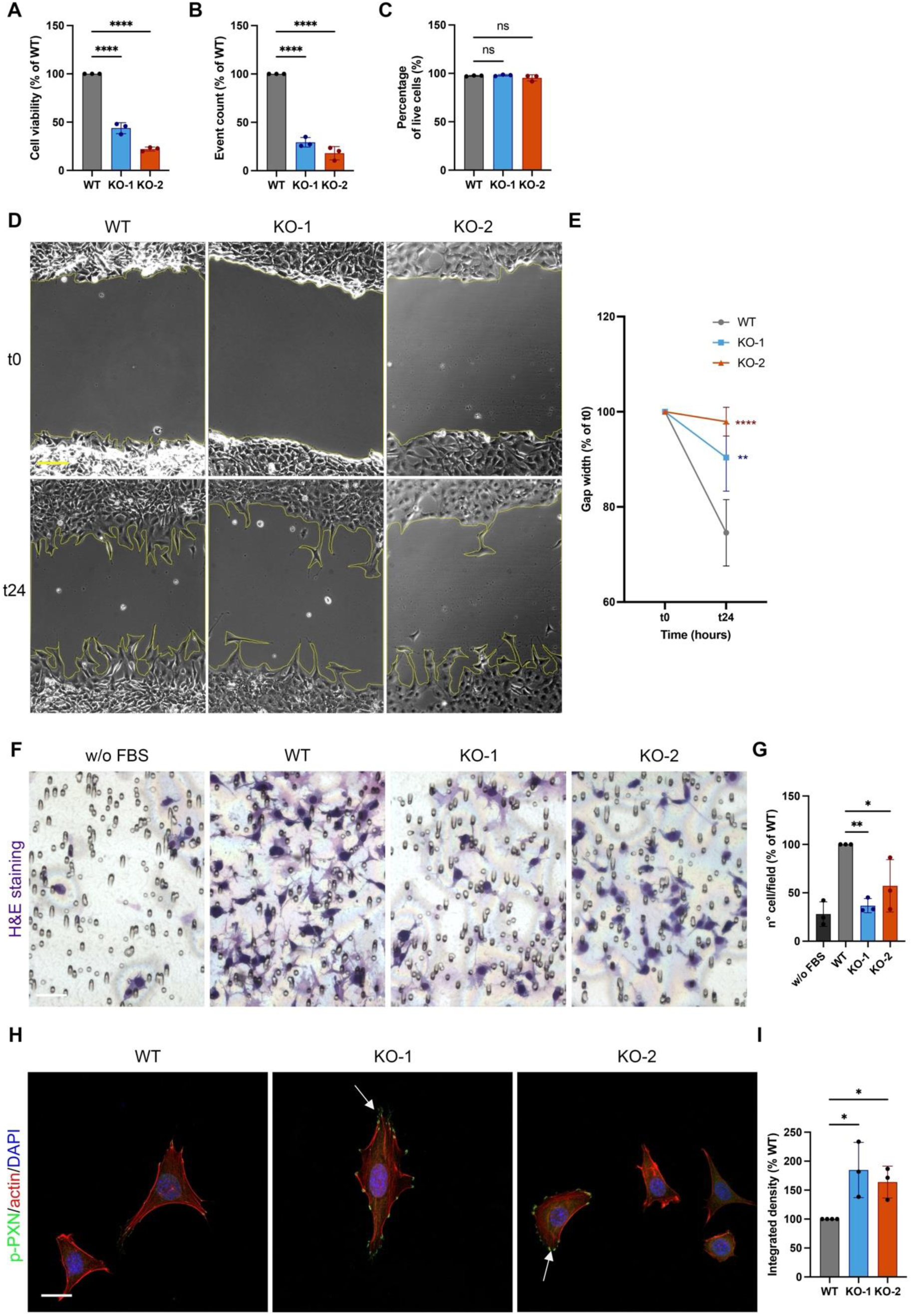
Phenotypic characterization of *Chd7*-KO GN11 cells. (A) Cell viability analysis by MTT assay performed at 72 h after seeding in WT and *Chd7*-KO GN11 cells (KO-1 and KO-2). Data are expressed as % of WT and shown as mean ± SD (N = 3; One-way ANOVA followed by Dunnett’s multiple comparisons test, **** p < 0.0001). (B) Flow cytometry-based quantification of total cellular events at 72 h after seeding, expressed as % of WT GN11 cells. Data are shown as mean ± SD (N = 3; One-way ANOVA followed by Dunnett’s multiple comparisons test, **** p < 0.0001). (C) Flow cytometry analysis of cell viability in WT and *Chd7*-KO cells using LIVE/DEAD staining at 72 h after seeding. Data are presented as mean ± SD (N = 3; ns, non significant, WT vs KO-1 p = 0.9140, WT vs KO-2 p = 0.3587). (D,E) Analysis of GN11 WT and *Chd7*-KO cell migration by wound healing assay performed under serum-free conditions (without FBS). (D) Representative phase-contrast images of scratch wounds at the time of scratching (t0) and after 24 h (t24). Yellow lines indicate wound edges. Scale bar: 500 μm. (E) Quantification of wound gap width over time, expressed as % of initial gap at t0. Data are shown as mean ± SD (N = 3; two-way ANOVA followed by Dunnett’s multiple comparisons test; ** p = 0.0012, **** p < 0.0001). (F,G) Assessment of chemotactic migration in GN11 WT and *Chd7*-KO cells towards using transwell assays toward a chemoattractant (FBS). (F) Representative bright-field images of migrated WT and *Chd7*-KO cells on the lower surface of the membrane. Scale bar: 250 μm. (G) Quantification of of migrated cells per field, expressed as % of WT GN11 cells. Data are shown as mean ± SD (N = 3; one-way ANOVA followed by Dunnett’s multiple comparison test, * p = 0.0310, ** p = 0.0054). (H,I) Analysis of focal adhesion components in WT and *Chd7*-KO GN11 cells by immunofluorescence staining for phosphorylated paxillin (p-PXN, green), F-actin (red), and nuclei (DAPI, blue). (H) Representative confocal images Scale bar: 20 μm. (I) Quantification of integrated p-PXN fluorescence intensity. Data are shown as mean ± SD (N = 3; one-way ANOVA; WT vs KO-1 * p = 0.0129, WT vs KO-2 * p = 0.0452). Abbreviations: H&E, haematoxylin and eosin, FBS, foetal bovine serum; p-PXN, phosphor-paxillin.

To determine whether the reduced cellular metabolic activity reflected increased cell death or a slower proliferation rate, consistent with the known role of CHD7 (Ohta *et al*, 2016; Yao *et al*, 2020; Layman *et al*, 2009), we first quantified cell numbers by flow cytometry at the same time points used for the MTT assays. KO clones displayed a significant decrease in event counts compared to WT cells at 24, 48 (**Fig. EV2C**) and 72 h (**Fig. 3B**), indicating impaired proliferative capacity. Next, we assessed cell death by flow cytometry using LIVE/DEAD staining. No significant differences in the percentage (%) of live cells were detected between KO and WT cells 72 h after seeding (**Fig. 3C**), confirming that the observed phenotype arises from diminished proliferation rather than increased apoptosis.

In parallel, given that transcriptomic analyses revealed dysregulation of pathways involved in cell migration and that CHD7 has been previously shown to play a crucial role in cell migration across different contexts (Schulz *et al*, 2014; Sanosaka *et al*, 2022; Okuno *et al*, 2017), we determined whether *Chd7* loss similarly impacts the migratory behaviour of immature GnRH neurons by migration assays.

First, we performed wound healing assays under serum-starved conditions, to specifically assess migration minimizing proliferation-driven effects on gap closure. Under these conditions, WT GN11 cells demonstrated a time-dependent closure of the scratch gap over 24 h, reflecting their intrinsic migratory capacity. In contrast, both *Chd7* KO-1 and KO-2 clones showed a reduced ability to close such gap (**Fig. 3D**), with a significantly wider gap compared to WT cells (**Fig. 3E**). To further validate this migratory impairment, we performed transwell migration assays. Consistent with wound healing assays, both KO-1 and KO-2 clones exhibited a significant reduction in chemotactic migration towards foetal bovine serum (FBS) compared to WT cells **(Fig. 3F,G**), indicating that *Chd7* loss impairs both chemokinesis and chemotaxis, ultimately compromising the overall migratory competence of GN11 cells.

Given the intimate interplay between cell migration and adhesion, particularly through focal adhesion complexes, we next examined whether the loss of *Chd7* could influence cell adhesion dynamics. Focal adhesions are regulated by key signalling molecules, including paxillin (PXN), a scaffolding protein involved in linking integrin-mediated adhesion to the actin cytoskeleton (Rashid *et al*, 2017). The phosphorylated form of paxillin (p-PXN) serves as a marker of active focal adhesions and is associated with enhanced cell-substrate adhesion and reduced motility (Deakin & Turner, 2008; Lettieri *et al*, 2019). To assess changes in focal adhesion signalling, we performed immunofluorescence experiments using an antibody specific for p-PXN in WT and *Chd7*-KO GN11 cells. Notably, KO clones displayed an increased number of p-PXN^+^ *punctae* at focal adhesion contacts compared to WT cells (**Fig. 3H**). Quantification of the fluorescent signal intensity, by integrated density, confirmed a statistically significant increase of p-PXN signal in KO cells (**Fig. 3I**), suggesting enhanced focal adhesion activity.

Overall, these data demonstrate that *Chd7* deficiency compromises both the proliferative capacity and the migratory competence in immature GnRH neurons, consistent with the transcriptomic profile of KO clones.

### *chd-7(gk290) C. elegans* mutants exhibit fertility defects

To rapidly identify possible drugs able to bypass *Chd7* deficiency, we took advantage of *C. elegans,* an invertebrate model allowing rapid and large drug screenings (Artal-Sanz *et al*, 2006). We selected the *chd-7(gk290)* allele to be tested in the drug screening in *C. elegans,* as being the most used and characterized (Schmeisser *et al*, 2017; Jamadagni *et al*, 2021; Rawsthorne *et al*, 2021; Jofré *et al*, 2022). Since it has been reported that other *chd-7* alleles show altered gonad proliferation and migration (Jofré *et al*, 2022) and that fertility-related phenotypes were observed in *Sema* gene mutants (Dalpé *et al*, 2005; Liu *et al*, 2005), we investigated the fertility of *chd-7(gk290)* mutants. We first observed that mutant animals exhibited a “bag-of-worms” phenotype (Trent *et al*, 1983), caused by egg retention and hatching within the mother’s body, which might be due to a vulva defect during development. The penetrance of this defect increased with time and was observed in about 30% of *chd-7* mutant worms six days (144 h) post-hatching (**Fig. 4A**). Such an alteration is correlated to a defect in egg-laying (Trent *et al*, 1983), thus we counted the number of eggs laid by adult *chd-7* mutants and WT every day for four days. Compared to WT animals, *chd-7* mutant animals laid fewer eggs, a phenotype already visible after 48 h of observation (**Fig. 4B**). This difference was more pronounced after 72 and 96 h while after 120 h both WT and mutant animals had almost stopped laying eggs. Our findings reveal that *chd-7*(*gk290*) mutant animals exhibit a defect in fertility and lay in total approximately half as many eggs compared to WT animals (**Fig. 4C**). This interesting and tractable phenotype is ideal for drug screening, as it requires a short developmental time (three days) and only simple manipulations, making it well suited for a semi-automated screening approach.

**Figure 4.**
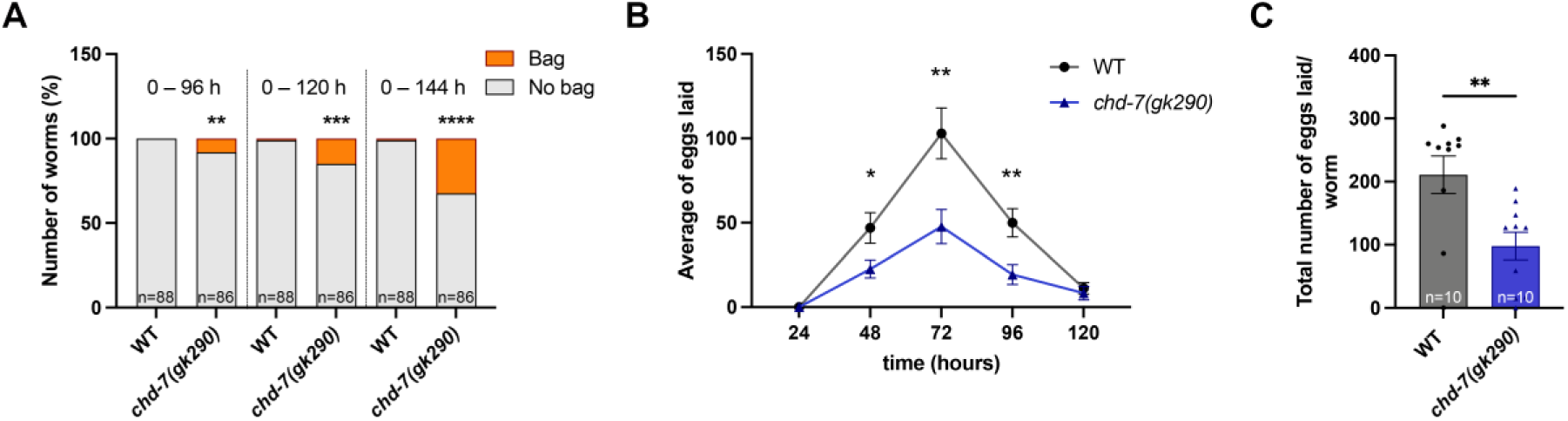
*chd-7(gk290)* animals show a defect in egg-laying. (A) Percentage of WT or *chd-7(gk290)* animals with normal (No bag, grey) or bag-of-worms phenotype (Bag, orange) at different time intervals. 0 refers to the day when animals start hatching. Compare two proportions non-parametric z-test: ** p = 0.0066, *** p = 0.0007, **** p < 0.0001. Number of worms analysed (n) per time point: WT, n = 88; *chd-7(gk290)* n = 86. (B) Average number of eggs laid by a single animal during time by the two different genotypes: black line WT; blue line *chd-7(gk290)*. Hours are referred to the time passed after the L4 larval stage. Each dot is the mean of the eggs laid by 10 animals (n = 10) and error bars are SEM. Unpaired t-test: * p = 0.0330, ** p < 0.0065. (C) Average number of total eggs laid by 10 animals (dots) in 96 h after the L4 larval stage by WT (grey bar and circles) or *chd-7(gk290)* animals (blue bar and triangles). Bars are the mean and error bars are SEM. Unpaired Mann-Whitney test: ** p = 0.002.

### Semi-automated screening of an epi-drug compound library in *C. elegans*

To establish a quantitative platform for phenotype-based screening, we exploited the egg-laying defect of *chd-7(gk290)* mutants in a liquid multi-well format. Animals were grown in liquid medium supplemented with bacteria as source of feeding, and the absorbance of each well was measured at 595 nm every day, using a plate reader (**Fig. 5A**). The screening is based on the fact that at time zero only eggs are present in wells, leading to high well turbidity due to high bacterial concentration. As the experiment progresses, these initial eggs hatch and develop into larvae which eat bacteria. Once these animals reach adulthood, they begin to lay new eggs, which subsequently hatch and rapidly expand the population. The difference in the turbidity measured after 96 hours is driven by the total number of animals present in the well: the consumption of bacteria is carried out by both the original adults and their numerous hatched offspring. Since WT animals lay twice as many eggs as *chd-7* mutants, they produce a significantly larger total population over the 96-hour period. Consequently, the bacterial consumption by this larger group leads to a much lower turbidity and absorbance in WT wells compared to *chd-7(gk290)* wells, where a smaller population leaves the medium noticeably more turbid (**Fig. 5B**). With this setup we screened a library of 234 chromatin modifiers, and we have successfully identified molecules able to reduce the turbidity of wells containing *chd-7(gk290)* animals compared to the mock (e.g., molecules 17 and 20 in **Fig. 5C**). Therefore, these molecules are probably able to partially rescue the defect of deposition. Moreover, we also identified molecules that increase the turbidity compared to the mock (e.g., molecules 1 and 4 in **Fig. 5C**) and that probably have a toxic effect on animals or worsen the deposition defect. A heatmap schematizing the effect of all the 234 compounds tested was prepared by calculating the difference (Δ) in turbidity with mock treated animals (**Fig. EV3A)**. We arbitrarily identified a Δ-turbidity threshold to select a broad range of promising compounds for further characterization and avoid losing potentially interesting molecules. Thus, we selected 72 molecules causing a decrease in turbidity (5% < Δ-turbidity values < 100%) and 30 molecules increasing turbidity (-15% > Δ-turbidity values > -100%) as positive hits among the 234 compounds initially screened (Discovery step) (**Appendix Table S1**).

**Figure 5.**
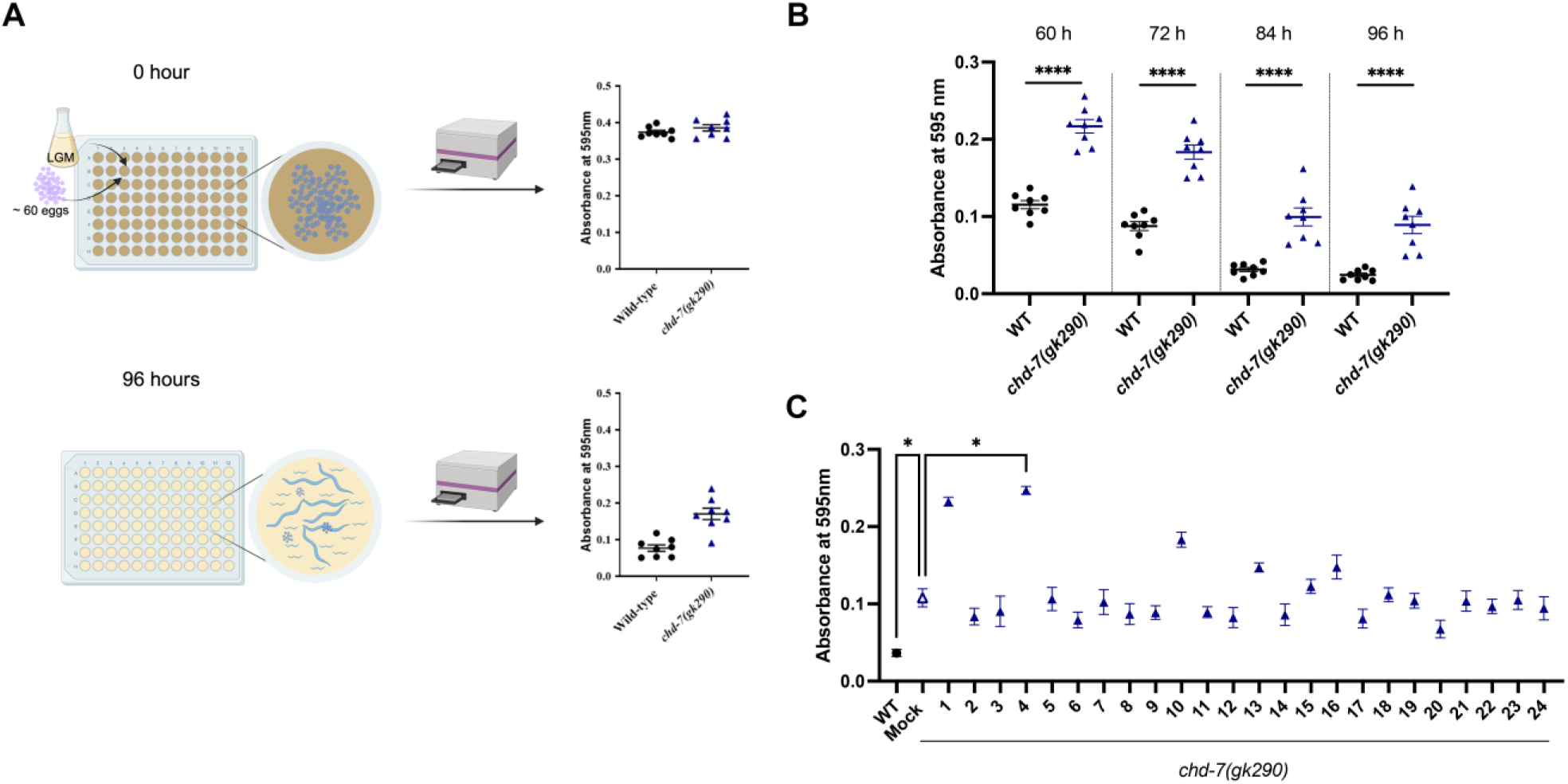
Screening of an epi-drug compound library. (A) Visual scheme representing the set-up of the *C. elegans* screening. At time 0 h wells were seeded with ∼ 60 eggs of WT or *chd-7(gk290)* animals in Liquid Growth Media (LGM). Absorbance at 595 nm was immediately measured revealing high turbidity due to elevated bacterial concentration. After 96 h (end point of the screening) both strains reached adulthood and began to lay eggs, which subsequently hatch and rapidly expand the population. WT animals produced twice as many eggs as *chd-7* mutants. As a result, the higher population density of WT animals led to increased bacterial consumption and a more consistent reduction in well turbidity compared to mutant-containing wells. Created with BioRender.com. (B) Absorbance at 595 nm of eight wells per condition after growing 60 eggs of WT (black circles) and *chd-7(gk290)* (blue triangles) at different time points from starting the experiment. 0 refers to the day when animals start hatching and the treatment started. Each dot represents the absorbance of a single well (n = 8 per genotype at each timepoint). Mean and SEM are shown. Unpaired t-test, **** p < 0.0001. (C) Example of the average absorbance at 595 nm of the eight wells filled with 24 compounds from the epi-drug library after 96 h of treatment: black circle represents WT animals treated with mock; empty blue triangle is *chd-7(gk290)* animals treated with mock; filled blue triangles are *chd-7(gk290)* animals treated with 24 different compounds. Each dot represents the absorbance of the average of the eight replicate wells for each condition. Mock treatment is DMSO 1%. Error bars are SEM. One-way ANOVA with Kruskal-Wallis test; Mock vs WT * p = 0.0440; Mock vs 4 * p = 0.0387.

### Validation of positive hits in *C. elegans*

To identify the most promising candidates, all the selected molecules from the discovery screening step underwent a validation screening using the same experimental setup and we confirmed most of the hits. A summary of the results from the validation screening is presented in **Fig. EV3B** and the full list of the results is given in **Appendix Table S2**. By applying a more stringent Δ-turbidity cut-off, we further selected the best drugs as the ones changing Δ-turbidity over 8% or below -25%. The validation screening step confirmed the efficacy of 30 turbidity-decreasing molecules (possibly rescuing the defect) among 72 tested and 24 turbidity-increasing molecules (possibly worsening the defect) among 30. Since the goal of the screening was to identify molecules with selective activity toward *chd-7(gk290)* mutants, the 54 top candidates were subsequently assessed in a WT background (WT test step) (**Appendix Table S3**). The screening setup mirrored the previous one, with the exception that we screened WT animals, while *chd-7(gk290)* mutant animals treated with mock were used as controls. Intriguingly, six molecules affecting *chd-7* mutants had no effect on WT animals. Among these, five improved the aberrant egg-laying phenotype of mutant worms (highlighted in green in **Fig. EV3C**), whereas one further exacerbated it (highlighted in red in **Fig. EV3C**). Additionally, three molecules were identified to have opposite effects in the two genetic backgrounds (highlighted in purple in **Fig. EV3C**). The impact of these nine candidate hits on turbidity across the discovery, validation, and WT screening steps is presented in **Fig. 6A–C**. Overall, the screening process yielded nine potential molecules presenting a different effect in the two genetic backgrounds (**Table 1**).

**Figure 6.**
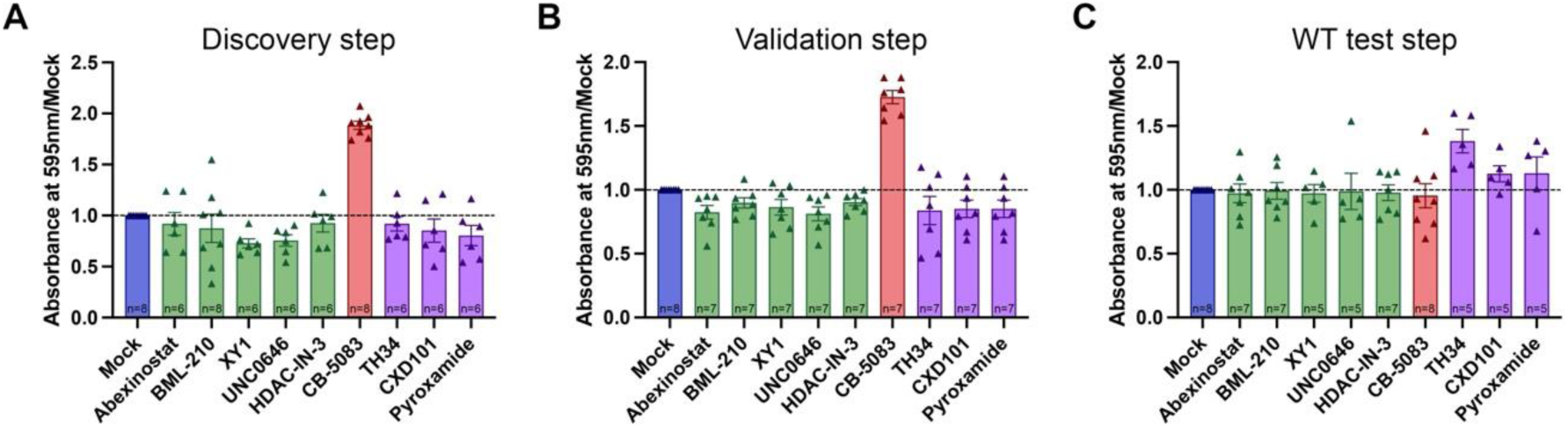
Impact of best candidate hits on bacterial turbidity across different screening stages. Absorbance values measured at 595 nm for the nine best candidate hits, normalized to the mock control (1% DMSO), during the discovery (A), validation (B), and WT test (C) screening steps. Green dots/bars represent molecules that induce a reduction in absorbance compared to mock treated animals (potential rescue), while the red dots/bars correspond to a molecule causing an increase in absorbance (phenotypic worsening). Molecules shown in purple exhibit divergent effects depending on the genetic background, since they reduce the absorbance in *chd-7* mutants (A and B) while increasing it in the WT background (C). Each dot represents the normalized absorbance value of a single well; n = number of wells analysed is reported in each bar. Error bars indicate SEM, dashed line is the mock control.

**Table 1.**
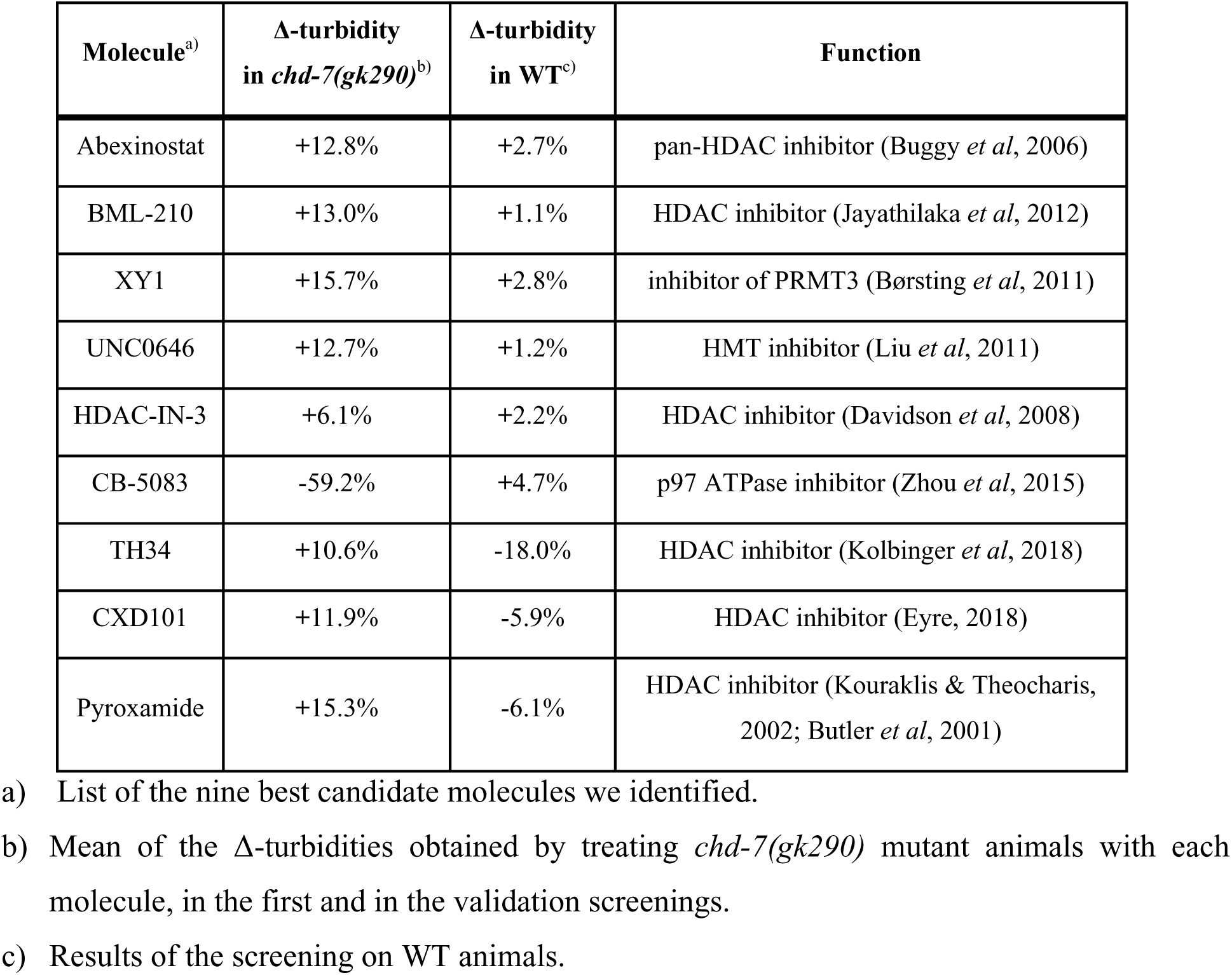
List of the best molecules identified in the semi-automated screening.

| Molecule<sup>a)</sup> | Δ-turbidity<br>in *chd-7(gk290)*<sup>b)</sup> | Δ-turbidity<br>in WT<sup>c)</sup> | Function |
| --- | --- | --- | --- |
| Abexinostat | +12.8% | +2.7% | pan-HDAC inhibitor (Buggy *et al*, 2006) |
| BML-210 | +13.0% | +1.1% | HDAC inhibitor (Jayathilaka *et al*, 2012) |
| XY1 | +15.7% | +2.8% | inhibitor of PRMT3 (Børsting *et al*, 2011) |
| UNC0646 | +12.7% | +1.2% | HMT inhibitor (Liu *et al*, 2011) |
| HDAC-IN-3 | +6.1% | +2.2% | HDAC inhibitor (Davidson *et al*, 2008) |
| CB-5083 | -59.2% | +4.7% | p97 ATPase inhibitor (Zhou *et al*, 2015) |
| TH34 | +10.6% | -18.0% | HDAC inhibitor (Kolbinger *et al*, 2018) |
| CXD101 | +11.9% | -5.9% | HDAC inhibitor (Eyre, 2018) |
| Pyroxamide | +15.3% | -6.1% | HDAC inhibitor (Kouraklis & Theocharis,<br>2002; Butler *et al*, 2001) |

a) List of the nine best candidate molecules we identified.
b) Mean of the $\Delta$ -turbidities obtained by treating *chd-7(gk290)* mutant animals with each molecule, in the first and in the validation screenings.
c) Results of the screening on WT animals.

### Validation of best hits in *Chd7*-deficient cells reveals phenotypic rescue of proliferation and migration

Following the identification of candidate molecules in the *C. elegans chd-7* mutant model, we next sought to evaluate their therapeutic potential in a mammalian system. To this end, we employed our *Chd7*-KO murine neuronal model to investigate the ability of selected compounds to rescue the previously observed defective cellular phenotypes. We focused on the five candidate compounds that significantly rescued the mutant phenotype without affecting WT animals.

First, we performed MTT assays to assess compound cytotoxicity and to identify an effective concentration range for downstream experiments. To this end, WT and *Chd7*-KO-1 (representative of both KO clones) cells were treated with increasing doses of each compound (10 nM-100 µM) for 24 and 48 h and viability was assessed normalized to vehicle-treated controls (0.5% DMSO). As shown in **Fig. EV4**, all compounds showed dose-dependent effects. Concentrations ≤1 µM showed minimal cytotoxicity, whereas higher doses (50-100 µM) of some compounds significantly reduced cell viability, particularly in WT cells. Based on these results, we selected the highest non-cytotoxic concentration for each compound after 48 h of exposure. Specifically, we chose doses that maintained viability in both KO and WT backgrounds, thereby ensuring that the selected concentrations were not generally toxic to GN11 cells. The concentrations used in all subsequent assays were: 100 nM for Abexinostat, 1 μM for BML-210, 50 μM for XY1, 1 μM for UNC0646, and 100 nM for HDAC-IN-3.

We next evaluated whether the candidate compounds could rescue the proliferation defects associated with *Chd7* loss. Specifically, we employed automated High-Content Screening (HCS) of DAPI-stained nuclei 72 h after seeding to confirm a significant reduction in cell numbers in both KO clones compared to WT GN11 cells (**Fig. 7A**), validating previous results (**Fig. 3A,B**). This platform served as a fast and reproducible tool for subsequent high-throughput assays using proliferative capacity as phenotypic readout. HCS of KO and WT cells following a 48-hour drug treatment revealed that only XY1 significantly increased proliferation in both KO-1 and KO-2 cells relative to vehicle-treated controls (NT), while having no effect on WT cells (**Fig. 7B-D**). In contrast, BML-210 significantly reduced proliferation in KO-1 and WT cells, indicating a non-specific growth-inhibitory effect. Abexinostat selectively decreased proliferation in WT cells but not in either KO line. UNC0646 and HDAC-IN-3 did not significantly alter proliferation in any genotype.

**Figure 7.**
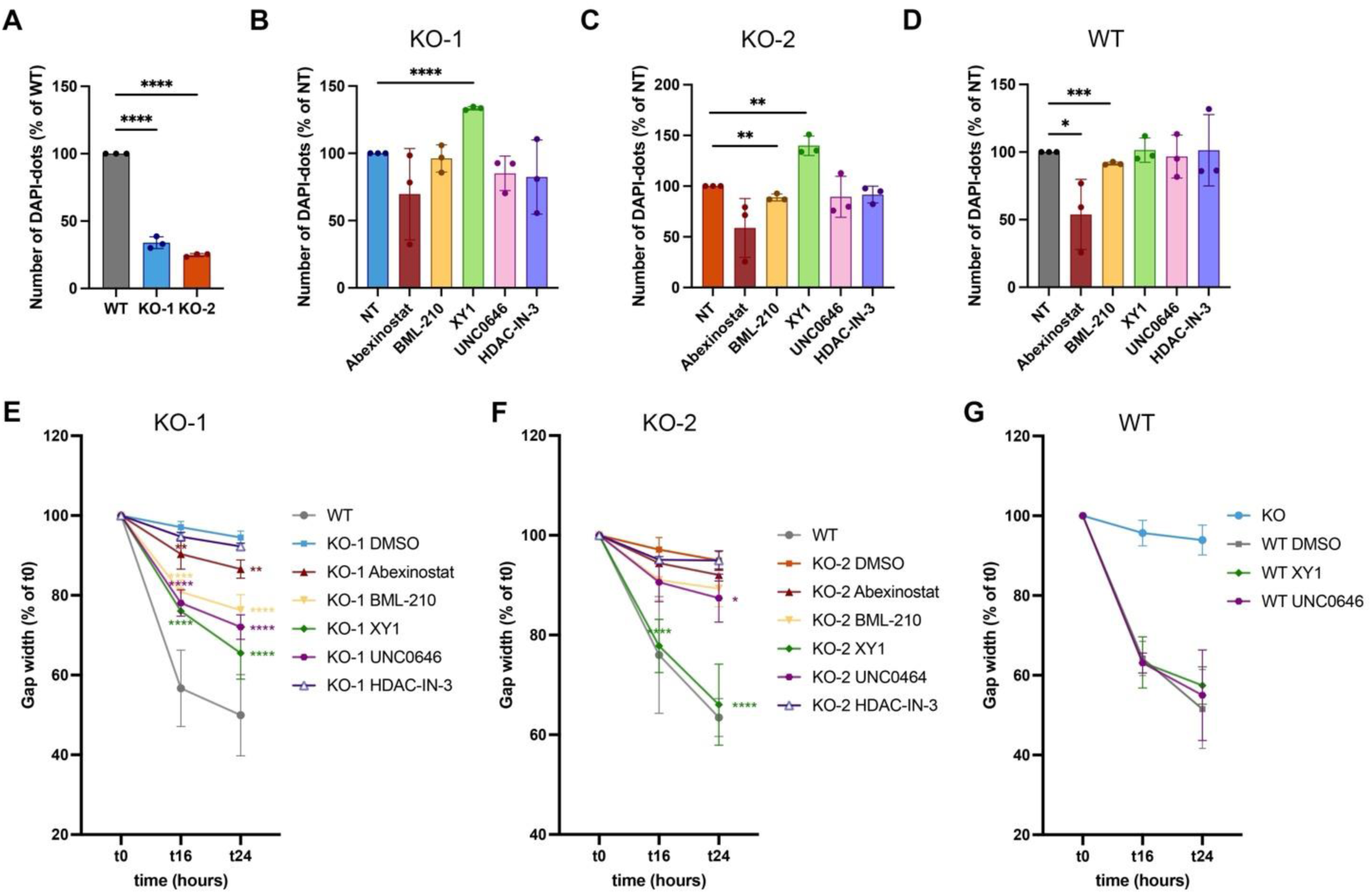
Pharmacological modulation of proliferation and migration in *Chd7*-KO GN11cells. (A) HCS-based quantification of DAPI-positive nuclei in WT and *Chd7*-KO GN11 cells (KO-1 and KO-2) at 72 h after seeding. Data are expressed as % of WT and shown as mean ± SD (N = 3; One-way ANOVA followed by Dunnett’s multiple comparisons test, **** p < 0.0001). (B–D) Quantification of DAPI-positive nuclei by HCS, expressed as % of vehicle-treated controls (NT, 0.5% DMSO), as a readout of cell proliferation following 48 h drug treatment with candidate compounds at selected non-toxic concentrations (Abexinostat, 100 nM; BML-210, 1 μM; XY1, 50 μM; UNC0646, 1 μM; HDAC-IN-3, 100 nM). Analyses were performed at 72 h after seeding *Chd7*-KO clones KO-1 (B) and KO-2 (C), and in WT GN11 cells (D). Data are shown as mean ± SD (N = 3; Unpaired t test, (B) **** p < 0.0001; (C) BML-210 vs NT ** p = 0.0048, XY1 vs NT ** p = 0.0020; (D) Abexinostat vs NT * p = 0.0371, BML-210 vs NT *** p = 0.0002). (E–G) Wound healing assays performed under serum-free conditions in KO-1 (E), KO-2 (F) and WT (G) GN11 cells. Graphs gap width was measured at 0, 16 and 24 h after scratch and expressed as % of the initial gap (t0). Cells were treated with candidate compounds immediately after scratch. Data are shown as mean ± SD (N = 3; 2-way ANOVA followed by Dunnett’s multiple comparison test; (E, t16) KO-1 DMSO vs KO-1 Abexinostat ** p = 0.0096, KO-1 DMSO vs BML-210, XY1 and UNC0646 **** p < 0.0001; (E, t24) KO-1 DMSO vs KO-1 Abexinostat ** p = 0.0020, KO-1 DMSO vs BML-210, XY1 and UNC0646 **** p < 0.0001; (F, t16) KO-2 DMSO vs XY1 **** p < 0.0001; (F, t24) KO-2 DMSO XY1 **** p < 0.0001, KO-2 DMSO vs UNC0646 * p = 0.02392). Abbreviations: NT, non-treated; DMSO, dimethyl sulfoxide.

We next assessed the impact of the selected compounds on cell migration performing wound healing assays in KO-1, KO-2 and WT GN11 neuronal cells (**Fig. 7E-G**). Cells were treated immediately after scratching and maintained under starvation conditions throughout the experiment. Wound closure was quantified relative to DMSO-treated controls for each genotype, after confirming that 0.5% DMSO had no significant effect on migration compared with untreated cells (**Fig. EV5**). WT GN11 cells served as positive controls for cell migration, while KO-1 cells were used as negative controls in the assays assessing the effects of candidate compounds on WT cells. As shown in **Fig. 7E**, treatment with Abexinostat, BML-210, XY1 and UNC0646 significantly increased wound closure over time in KO-1 cells, demonstrating a rescue of the *Chd7*-associated migration defect. This rescue effect was confirmed in KO-2 cells for XY1 and UNC0646 (**Fig. 7F**). Notably, none of the tested compounds altered the migration of WT cells (**Fig. 7G**), supporting a genotype-specific rescue. In contrast, HDAC-IN-3 failed to improve migration in either KO clone (**Fig. 7E,F**).

Overall, these findings demonstrate that XY1 and UNC0646 selectively rescue the proliferation and migration defects caused by *Chd7* loss, supporting their potential as genotype-specific modulators of Chd7-dependent cellular phenotypes.

### XY1 and UNC0646 treatment partially restores aberrant semaphorin gene expression in *Chd7*-depleted GN11 cells

In our murine CS model, bulk RNA-seq revealed consistent upregulation of SEMA family genes (*Sema3a*, *Sema3f*, *Sema3b*) and plexin receptors (*Plxnd1*, *Plxnb1*) (**Fig. 2E,F**) as a downstream consequence of *Chd7* loss. We therefore investigated whether selected epigenetic compounds could restore the expression of these genes in vitro.

Based on their ability to rescue both proliferation and migration defects, XY1 and UNC0646 were selected for targeted gene expression analyses after 48 h treatment. As shown in **Fig. 8A**, both compounds significantly reduced *Sema3a* expression in KO-1 and KO-2 cells, reversing the upregulation caused by *Chd7* depletion. Additional effects showed greater variability, with UNC0646 decreasing *Sema3f* expression in KO-2 cells (**Fig. 8B**), while reducing *Plxnd1* expression only in KO-1 cells (**Fig. 8C**).

**Figure 8.**
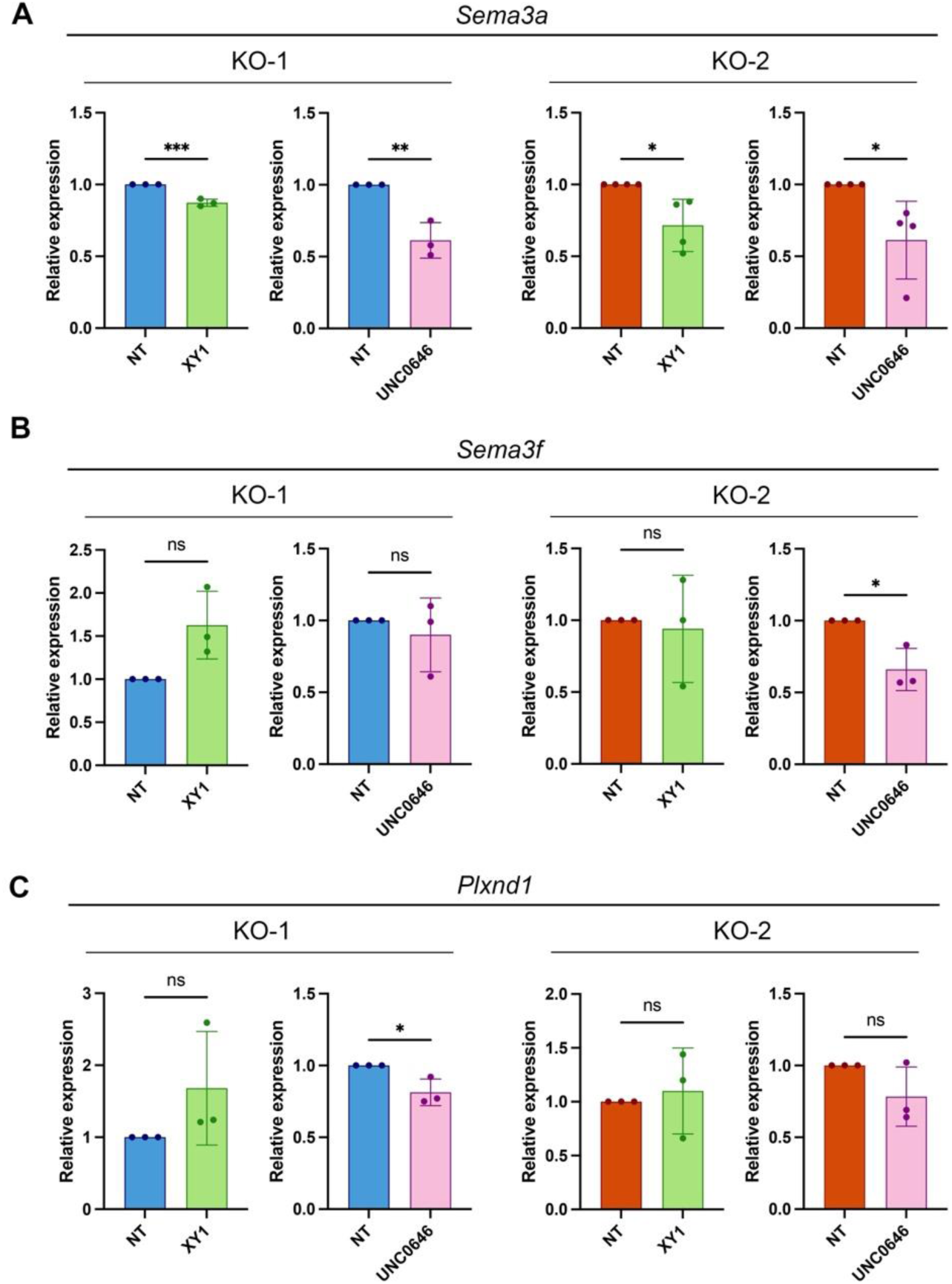
Modulation of semaphorin and plexin gene expression in *Chd7*-KO GN11 cells by selected epidrugs. qPCR analysis of indicated *Sema* and *Plxn* transcripts in *Chd7*-KO GN11 cell clones KO-1 (light blue) and KO-2 (orange) following 48 h with XY1 (50 µM, green) or UNC0646 (1 µM, pink). Transcript levels were normalized to *Gapdh* and expressed relative to untreated (NT; vehicle-only, 0.5% DMSO) KO controls. Data are shown as mean ± SD (N = 3/4; Unpaired t-test, *p < 0.05, **p < 0.01, ***p<0.001, ns, not significant; specific p values are reported in **Appendix Table S4**).

These results indicate that XY1 and UNC0646 can partially normalize the SEMA-related transcriptional alterations induced by *Chd7* loss.

## Discussion

This study outlines a multi-model, phenotype-driven pharmacological screening strategy to identify novel therapeutic candidates for CS, with a specific focus on targeting SEMA signalling as downstream to CHD7. Our findings provide the first experimental evidence that pharmacological epigenetic modulation can rescue neurodevelopmental and neuroendocrine defects caused by *CHD7* haploinsufficiency.

Semaphorins, together with their receptors, neuropilins and plexins, collectively referred to here as SEMA genes, were originally identified as axon guidance cues. However, they are now recognized as key regulators of central nervous system (CNS) patterning during embryonic development, orchestrating critical processes such as neuronal proliferation, differentiation, and migration (Oleari *et al*, 2019; Alto & Terman, 2017). Variants in *SEMA3A* and *SEMA3E*, both crucial for neuronal migration and survival (Cariboni *et al*, 2011b, 2015), are associated with multiple neurodevelopmental disorders, including CS, thus suggesting a functional interplay with CHD7 (Lalani *et al*, 2004; Cariboni *et al*, 2015; Hanchate *et al*, 2012; Ufartes *et al*, 2018; Schulz *et al*, 2014). In line with this, genome-wide studies in mouse and human neural stem cells (NSCs) have identified CHD7 binding at active enhancer regions of several *SEMA* genes, including *SEMA3A* and *SEMA3F* (Chai *et al*, 2018; Engelen *et al*, 2011). Consistently, *CHD7* depletion has been shown to dysregulate *SEMA* gene expression in multiple vertebrate models, including mouse, zebrafish, and *Xenopus laevis* (Ufartes *et al*, 2018; Schulz *et al*, 2014; Liu *et al*, 2019). Despite these associations, no prior studies have directly explored the therapeutic potential of targeting SEMA pathways in CS.

Using both a CRISPR/Cas9-engineered *Chd7*-KO murine neuronal line and a *C. elegans chd-7* mutant strain, we established a mechanistic link between *Chd7* loss, SEMA gene dysregulation, and CS-relevant phenotypes. The *Chd7*-KO GN11 model mirrors key cellular phenotypes associated to CS, including reduced proliferation and impaired migration (Yao *et al*, 2020; Okuno *et al*, 2017). Transcriptomic profiling revealed that *Chd7* loss disrupted pathways related to cell cycle, migration and adhesion, with consistent upregulation of *SEMA* gene, particularly *Sema3a*. Elevated *Sema3a*, a known repulsive guidance cue for GnRH neurons (Cariboni *et al*, 2011b), likely contributes to impaired migration, supporting its role as a pathogenic driver and suggesting a pathological feedback loop in which aberrant repulsive signalling may exacerbate migration defects due to *Chd7* loss. This observation aligns with and extends previous findings linking CHD7 to SEMA gene regulation and GnRH neuron development (Lettieri *et al*, 2021). Notably, our transcriptomic data contrast with earlier studies reporting CHD7 as a positive regulator of *Sema3a* (Ufartes *et al*, 2018; Schulz *et al*, 2014; Feng *et al*, 2017a; Schnetz *et al*, 2010), suggesting that CHD7 regulatory effects on SEMA genes may be strongly context- and stage-dependent during development.

To complement our mammalian studies, we leveraged *C. elegans*, where the highly conserved *chd-7* gene offers a genetically tractable platform for functional genomics and drug screening (van Ravenswaaij-Arts & Martin, 2017). *C. elegans* has emerged as a promising tool for compound screening assays, through the development of rapid worm-based screening methods that involve the cultivation of a large number of worms, which can be easily grown in liquid and in multi-well formats (Artal-Sanz *et al*, 2006; Giunti *et al*, 2021). Moreover, as observed in humans, also in *C. elegans* distinct mutations in *chd-7* cause different symptoms, with a certain variability of severity (Wong *et al*, 2019). The SEMA pathway genes are also conserved in *C. elegans* and their role in development very well characterized (Nukazuka & Takagi, 2017; Bonner & O’Connor, 2011; Chisholm *et al*, 2016). Interestingly, combined *chd-7* and semaphorin deletion or overexpressing mutants share similar phenotypes affecting fertility and development (Dalpé *et al*, 2005, 2004; Pellegrino *et al*, 2011; Liu *et al*, 2005). Indeed, we demonstrated that *gk290* deletion cause a fertility defect possibly due to vulva defects that resemble the ones observed in semaphorin pathway gene mutants, strongly supporting for a conserved genetic interaction. Thus, *C. elegans* offers the unique opportunity to study CHD7 and SEMA interactions in vivo and to perform unbiased drug screenings.

Indeed, in this study, thanks to the advantages of using *C. elegans*, a screening system was developed to identify new therapeutic targets for CS using the abnormal fertility phenotype of *chd-7(gk290)* animals. This system enabled a drug screening of 234 chromatin-modifying compounds. Starting with 234 compounds, this setup reduced the number of positive hits to 9. Interestingly, the entire screening process, was completed in approximately a month and a half, once again highlighting the great potential of *C. elegans* for the identification of new drugs in a short time. Moreover, among the positive hits, we identified nine hits targeting histone deacetylase (HDAC), protein arginine methyltransferase (PRMT), and histone methyltransferase (HMT) enzymes. Five of these compounds selectively decreased the bacterial turbidity observed when cultivating *chd-7* mutants, suggesting a rescue in fertility defects and highlighting the importance of epigenetic regulation in CS. Notably, these hits were distinct from those identified in another drug screening performed in *C. elegans* on a different mutant allele and phenotype (Jamadagni *et al*, 2021), underscoring the specificity and robustness of our strategy.

Two compounds, XY1 and UNC0646, were further validated in *Chd7*-KO GN11 cells, where they restored proliferation and migration defects and, more importantly, normalized *Sema3a* expression. Although their effects were partially clone-dependent, the significant downregulation of *Sema3a* by both compounds support for the therapeutic relevance of using these drugs to modulate SEMA signalling and bypass *CHD7* haploinsufficiency. UNC0646 exhibited broader transcriptional effects restoring other SEMA gene expression, suggesting a more comprehensive mechanism of action. However, its efficacy also seems partially dependent on genetic background or clonal variability, highlighting the need for further validation in additional models. UNC0646 functions as an HMT inhibitor, implicating epigenetic remodelling by this class of histone-modifying enzymes as a central mechanism in CS pathogenesis. This aligns with recent findings demonstrating that CHD7 activates neural transcriptional programs by removing the repressive histone mark H3K27me3 and enhancing chromatin accessibility. In *CHD7*-deficient neuroepithelial cells and human forebrain organoids, the loss of *CHD7* reduces both the proliferation and the differentiation of neural progenitors, resulting in CS-like brain anomalies such as microcephaly and olfactory bulb agenesis (Huang *et al*, 2025). Importantly, this study showed that inhibition of EZH2, the H3K27me3 methyltransferase, ameliorated neurodevelopmental defects in both human and murine *CHD7*-deficient models. This provides compelling evidence that targeting histone methylation can restore transcriptional balance and rescue developmental programs disrupted by *CHD7* loss. Given that UNC0646 similarly inhibits methyltransferases, its ability to downregulate *Sema3a* and broadly reprogram gene expression may reflect a parallel mechanism.

Future studies will be required to further elucidate the molecular targets and regulatory networks impacted by compounds such as XY1 and UNC0646 and investigate their efficacy, safety, and optimal timing of treatment during development in more complex vertebrate in vivo systems. Notably, the adaptability of this pipeline to screen alternative compound libraries, including FDA-approved drugs, could significantly accelerate the path from bench to bedside by enabling drug repurposing strategies. Such an expansion would increase the translational impact of our approach and help bridge the gap between experimental findings and clinical intervention in CS and potentially other neurodevelopmental disorders.

Taken together, our results support a model in which *Chd7* deficiency leads to aberrant upregulation of key axon guidance molecules, which may disrupt GnRH neuron migration and contribute to CS-associated neuroendocrine deficits. The ability of specific epigenetic modulators to selectively mitigate *Sema3a* dysregulation highlights a precise avenue for intervention, potentially addressing core pathogenic mechanisms underlying CS-associated HH and other neuroendocrine deficits.

Our multi-species platform thus establishes a powerful translational framework for therapeutic discovery in CS. By grounding compound selection in genetically defined, phenotypically validated models of *CHD7* deficiency, and targeting a pathway directly relevant to neuroendocrine development, this strategy offers a mechanistically precise alternative to prior efforts focused on broad chromatin modifiers or retinoic acid signalling (Huang *et al*, 2025; Micucci *et al*, 2013), and opens new avenues for the development of targeted therapies aimed at restoring reproductive and developmental function in CS.

Moreover, our work establishes a reliable and translatable screening platform that bridges findings from a *C. elegans* model to a mammalian neuronal context. The convergence of data across *C. elegans* and mammalian models strengthens the rationale for further preclinical investigation of compounds like XY1 and UNC0646.

## Materials and methods

### In vitro experiments

#### Cells

GN11 cells were cultured at 37°C in a humidified atmosphere containing 5% CO₂ in Dulbecco’s Modified Eagle’s Medium (DMEM; Euroclone, Milan, Italy; cat. ECB7501L), supplemented with 10% (v/v) heat-inactivated fetal bovine serum (FBS; Life Technologies, Monza, Italy; Cat. 10270106), 2 mM L-glutamine (Euroclone, Milan, Italy; cat. ECB3000), 100 IU/mL penicillin, and 100 µg/mL streptomycin (Euroclone, Milan, Italy; cat. ECB3001D). This medium is referred to as complete DMEM. Subconfluent cultures were dissociated with trypsin and reseeded in 57 cm² dishes at a density of 8 × 10⁴ cells/dish for routine passaging.

#### CRISPR/Cas9 genome editing

*Chd7*-deficient (*Chd7*-KO) GN11 cell clones were generated using the CRISPR/Cas9 system. Single-guide RNAs (sgRNAs) targeting exon 2 and exon 38 of the murine *Chd7* gene (GRCm38.p6, chromosome 4: NC_000070.6; 8,690,402–8,868,449) were designed using CHOPCHOP (https://chopchop.cbu.uib.no) and ATUM DNA 2.0 (https://www.atum.bio/eCommerce/cas9/input). Top-scoring sgRNAs were validated using the SyntheGo guide verification tool (https://design.synthego.com/#/validate). Selected sgRNA sequences (sgRNA1_exon 2: 5’-GCAGCAGCCTCGTTCGTTTA-3’, 5’-TAAACGAACGAGGCTGCTGC-3’; sgRNA2_exon 38: 5’-AGACGGTCAAACATTGACTC-3’, 5’-GAGTCAATGTTTGACCGTCT-3’) were cloned into the pSpCas9(BB)-2A-Puro V2.0 vector (PX459; Addgene, Watertown, MA, USA; cat. 62988) using BbsI sites for co-expression with SpCas9. Plasmid ligation was verified by Sanger sequencing (hU6-F, 5’-GAGGGCCTATTTCCCATGATT-3’). Plasmids were transfected into sub-confluent GN11 cells using Lipofectamine 3000 (Thermo Fisher Scientific, Segrate, Italy; cat. L3000015). Transfected cells were selected with 3 µg/mL puromycin (Merck, Milan, Italy; cat. P9620) for 48 h. Surviving cells were serially diluted into 96-well plates to isolate single-cell clones. Clones were expanded and screened for *Chd7* deletion by genomic PCR. The oligonucleotide sequences used for screening are the following: control, 5’-ACGGGCATCCTGAACTTTGA-3’ and 5’-GGGCACAATAGGCATCCTGA-3’; exon 3, 5’-CAGCTAGTGAAGAGCGACGA-3’ and 5’-CTGGAATCCCCTGCAGCAAT-3’; deletion, 5’-CGTGGGATTCCCGTCAAACA-3’ and 5’- GTTACTCGTCGTTTCCGGTG-3’. See deletion detection strategy in **Figure EV1A**. PCR-positive clones were verified by Sanger sequencing using a deletion-specific oligonucleotide (5’-CGTGGGATTCCCGTCAAACA-3’).

#### Immunoblotting

Protein extracts from WT and *Chd7*-KO GN11 cells were analyzed by Western blot. Cells were lysed in a buffer containing 150 mM NaCl, 50 mM Tris-HCl (pH 7.4), and 1% Triton X-100, supplemented with protease and phosphatase inhibitors (Roche, Monza, Italy; cat. 11697498001 and 4906837001). Lysates were clarified by centrifugation (13,000 rpm, 10 min, 4 °C) and protein concentration was determined using the Bradford assay (Bio-Rad Laboratories, Segrate, Italy, cat. 5000006). Proteins were resolved by SDS-PAGE and transferred to nitrocellulose membranes (Bio-Rad Laboratories, Segrate, Italy, cat. 1620112) by wet transfer overnight. Membranes were blocked in 5% non-fat dry milk in Phosphate Buffered Saline (PBS; Euroclone, Milan, Italy; cat. ECB4004L) 0.1% Tween-20 (Merck, Milan, Italy; cat. P1379-100ML) for 1 h and incubated with rabbit anti-CHD7 (D3F5; 1:500; Cell Signaling Technology, Leiden, Netherlands; cat. 6505) and anti-GAPDH (D16H11; 1:1000; Cell Signaling Technology, Leiden, Netherlands; cat. 5174T) antibodies, followed by HRP-conjugated anti-rabbit IgG (1:10,000; Santa Cruz Biotechnology, Dallas, TX, USA; cat. sc-2357-CM). Immunoreactive bands were visualized using enhanced chemiluminescence reagents (WASRAR ηC Ultra-2.0; Cyanagen, Bologna, Italy; cat. XLS075) according to the manufacturer’s protocol.

#### Immunocytochemistry and quantification

For immunostaining, WT and *Chd7*-KO cells were seeded on 13-mm glass coverslips in 24-well plates. For Chd7 staining, cells were plated at 5 × 10³ cells/well and fixed after 72 h; for phospho-paxillin (p-PXN) staining, cells were plated at 3 × 10⁴ cells/well and fixed after 3 h. Cells were fixed with 4% paraformaldehyde (PFA, Life Technologies, Monza, Italy; cat. 28908) for 15 min, permeabilized with 0.1% Triton X-100 in PBS (PBT), and blocked with 10% normal horse serum. Samples were incubated overnight at 4 °C with rabbit anti-CHD7 (1:100; Novus Biologicals, Bio-Techne Ltd., Abingdon, UK; cat. NBP1-77393SS) or rabbit anti-p-PXN (1:150; Cell Signaling Technology, Leiden, Netherlands; cat. 69363), followed by Alexa Fluor 488-conjugated anti-rabbit IgG (1:200; Jackson ImmunoResearch, Cambridgeshire, UK; cat. 111-545-003) for 2 h at room temperature. Nuclei were counterstained with DAPI (4’,6-diamidin-2-phenylindole, 1:10,000; Cell Signaling, Leiden, Netherlands; cat. 4083). For F-actin visualization, cells were stained with TRITC-conjugated phalloidin (1:400; Merck, Darmstadt, Germany; cat. P1951) for 30 min at 37 °C. Images were acquired using a Zeiss LSM900 confocal microscope equipped with Plan-Apochromat 40× (NA 1.3) and 63× (NA 1.4) objectives. Z-stacks were captured at 0.25 µm intervals and processed using ZEN 3.0 software (v.3.0.79.0006; Zeiss, Oberkochen, Germany). Quantification of p-PXN signal intensity was performed using ImageJ (v.1.54g; NIH). For each genotype, three independent experiments were performed and at least three images per experiment analyzed. Adobe Photoshop 2023 software (v. 24.0.0; Adobe, San Jose, CA, USA) was used to prepare the presented images.

#### RNA sequencing, data processing and visualization

Total RNA was extracted from frozen GN11 cell pellets with RNeasy Mini Kit (Qiagen, Hilden, Germany; cat. 74104) and RNA integrity assessed using Agilent 5300 Fragment Analyzer (Agilent, Palo Alto, CA, USA). Libraries were prepared using the Next Ultra II RNA Library Prep Kit for Illumina (New England Biolabs, Ipswich, MA, USA; cat. E7770). Validation and quantification of sequencing libraries were performed using NGS Kit on the Agilent 5300 Fragment Analyzer and Qubit 4.0 Fluorometer. Multiplexed libraries were sequenced using the Illumina NovaSeq 6000 instrument according to manufacturer’s instructions (Illumina, San Diego, CA, USA) with a 2x150 Pair-End (PE) configuration v.1.5. NovaSeq Control Software (v.1.7; Illumina, San Diego, CA, USA) was used for image analysis and base calling.

Raw sequencing reads were processed using Trimmomatic v.0.36 for adapter and quality trimming, and subsequently aligned to the *Mus musculus* reference genome (ENSEMBL, GRCm38.p6) using STAR v.2.5.2b. Gene-level read counts were obtained with featureCounts from the Subread package v.1.5.2, considering only uniquely mapped reads overlapping annotated exon regions. The resulting count matrix was used for downstream differential expression analysis. Differential gene expression between experimental groups was assessed using DESeq2. Statistical significance was determined using the Wald test, and genes with an adjusted *p-value* < 0.05 an absolute Log_2_ fold change (|Log2FC|) >1 were defined as differentially expressed genes (DEGs). Functional enrichment analysis of DEGs was performed using reString software (v.0.1.21; (Manzini *et al*, 2021)). Data visualization carried out as previously done using matplotlib and seaborn libraries for Python programming language (v.3.8.5) (Oleari *et al*, 2023; Amoruso *et al*, 2026; Hunter, 2007; Waskom *et al*).

#### MTT assay

Cell metabolic activity and drug cytotoxicity was evaluated using the MTT [3-(4,5-dimethylthiazol-2-yl)2,5-diphenyltetrazolium bromide] assay. Briefly, GN11 WT and *Chd7*-KO cells were seeded in 24-well plates at densities of 6 × 10⁴, 3 × 10⁴, or 1.5 × 10⁴ cells/well for assessment at 24, 48, and 72 h, respectively. At each time point, MTT (450 µg/mL in serum-free medium; Merck, Darmstadt, Germany; cat. M5655) was added and incubated for 30 min at 37 °C. After solubilization in isopropanol (Merck, Darmstadt, Germany; cat. 59300), absorbance was measured at 550 nm using an EnSpire Multimode Plate Reader (PerkinElmer, Shelton, CT, USA). For compound cytotoxicity assays, cells were treated with increasing concentrations (10 nM–100 µM) of each drug for 24 or 48 h. Vehicle-treated cells (0.5% DMSO) served as controls. Viability was normalized to vehicle controls and expressed as a percentage.

#### Flow cytometry

WT and *Chd7*-KO cells were seeded in 12-well plates at 1.5 × 10⁵, 7.5 × 10⁴, and 3.7 × 10⁴ cells/well to assess cell proliferation at 24, 48, and 72 h after medium change, respectively. At each time point, cells were harvested, centrifuged at 12,000 rpm for 5 min, washed once with PBS, centrifuged again, and mechanically disaggregated by pipetting to obtain a single-cell suspension in 100 μL PBS. Samples were analysed using the Novocyte3000 flow cytometer (Agilent, Palo Alto, CA, USA), and the number of cell events gated by size (Forward Scatter, FSC) and granularity (Side Scatter, SSC), was recorded as a measure of cell proliferation. Data were processed using NovoExpress software (v.1.4.1; Agilent, Palo Alto, CA, USA). For cell death assays, flow cytometry analysis was conducted 72 h after medium change using the Live/Dead Viability Assay kit (Thermo Fisher Scientific, Segrate, Italy; cat. L3224), according to the manufacturer’s instructions. For cell cycle analysis, cells were seeded in 12-well plates at a density of 1.5 × 10^5^ cells/well. After 6 h, cells were synchronized in G1 by serum starvation for 24 h, returned to complete medium, and collected 12 h later. Methanol-fixed cells were stained with FxCycle™ PI/RNase staining solution (Thermo Fisher Scientific, Segrate, Italy; cat. F10797) and analyzed by flow cytometry as described above.

#### High content imaging screening

WT and *Chd7*-KO GN11 cells were seeded in 48-well plates at a density of 2 × 10^3^ cells/well. For baseline proliferation analysis, cells were fixed 72 h after seeding. For proliferation analyses under drug treatment, cells were seeded at the same density and treated 24 h after plating with selected doses of each compound: 100 nM (Abexinostat, HDAC-IN-3), 1 μM (BML-210 and UNC0646), 50 μM (XY1). Cells were then fixed for analysis 48 h after treatment. Briefly, cells were fixed in 4% PFA for 15 min and permeabilized with PBT for 10 min at RT. Nuclei were counterstained with DAPI (1:10,000) and multiple images of each well were acquired using the ImageXpress Micro Confocal inverted microscope equipped with a 4× objective (Molecular Devices, San Josè, CA, USA). Automated quantification of DAPI-positive nuclei in each well was performed with the MetaXpress software (v. 6.6; Molecular Devices, San Josè, CA, USA).

#### Migration assays and quantification

Cell motility was evaluated using both wound-healing and transwell migration assays. For wound-healing assays, confluent GN11 WT and *Chd7*-KO monolayers were scratched and incubated in serum-free medium. For drug treatment, cells were exposed to non-toxic doses of candidate compounds in serum-free DMEM for 24 h. Phase-contrast images were captured at indicated time points using a Zeiss Axiovert 200 microscope equipped with a Photometric CoolSnap CCD camera (Roper Scientific, Ottobrun, Germany) and a Zeiss A-Plan 10× objective (NA 0.25). Wound closure was quantified in ImageJ software (v. 1.53a, Java 1.8.0_172; NIH, USA) relative to the initial wound area (time t0). Transwell migration assays were performed using 8 µm pore membranes (NeuroProbe, Gaithersburg, MD, USA; cat. PFB8). GN11 cells (2 × 10⁶ cells/mL) were allowed to migrate for 3 h at 37 °C toward complete DMEM (chemoattractant) or DMEM w/o FBS (negative control). Migrated cells were fixed, stained, and mounted onto glass slides as previously described (Cariboni *et al*, 2004). For quantitative analysis, membranes were observed using a Zeiss Axioskop 2 plus brightfield microscope and a Zeiss Plan-NEOFLUAR 20× objective (NA 0.5). Three randomfields of stained cells were counted for each well/condition, and the mean number of migrating cells/fields for each experimental condition was calculated. Adobe Photoshop 2023 software (v. 24.0.0) was used to prepare the presented images.

#### RNA extraction and Quantitative PCR (qPCR)

Total RNA was isolated from WT, *Chd7*-KO-1, and *Chd7*-KO-2 GN11 cells treated with XY1 (50 µM) or UNC0646 (1 µM) for 48 h using TriFast II Reagent (Euroclone, Milan, Italy; cat. EMR517100). RNA (1 µg) was reverse transcribed with the High-Capacity cDNA Reverse Transcription Kit (Thermo Fisher Scientific, Segrate, Italy; cat. 4368814). Quantitative PCR was performed using Luna Universal qPCR Master Mix (New England Biolabs, Ipswich, MA, USA; cat. M3003X) on a CFX96 Real-Time System (Bio-Rad Laboratories, Segrate, Italy), according to manufacturer indications and using specific oligonucleotides for murine *Sema3a* (5’-CGTCTTCCGGGAACCAACAA-3’, 5’-TGCACAGGCTTTGCCATAGA-3’), *Sema3f* (5’-AGACAGTGCCGTGGTTTCAA-3’, 5’-TCCTCTGCGCGAATCTCAC-3’), *Plxnd1* (5’-GGCCCACTACAAGATACCTGAG-3’, 5’-CATCCGTAGGTAGCACCAAATG-3’). Gene expression was normalized to *Gapdh* (5’-CATCCCAGAGCTGAACG-3’, 5’-CTGGTCCTCAGTGTAGCC-3’) using the ΔΔCt method.

### In vivo experiments

#### C. elegans strains

Nematodes have been grown and handled following standard procedures under uncrowded conditions on nematode growth medium (NGM) agar plates seeded with *Escherichia coli* strain OP50, unless otherwise noted (Brenner, 1974). NGM-Agar was prepared by dissolving 2.5 g peptone (Merck, Milan, Italy; cat. 70169), 3 g NaCl (Merck, cat. S3014), and 20 g agar (Merck, cat. A7002) in distilled water to a final volume of 1 L. The medium was autoclaved and cooled to approximately 65 °C before supplementation with 1 mL MgSO₄ 1 M (Merck, cat. 1.05886), 1 mL CaCl₂ 1 M (Merck, cat. 5080), 1 mL cholesterol in ethanol (5 mg/mL; Merck, cat. C3292), and 25 mL of 0.5 M phosphate buffer (pH 6.2). *C. elegans* strains used in this work were provided by the *Caenorhabditis* Genetic Center (CGC) funded by NIH Office of Research Infrastructure Programs (P40OD010440): the wild-type (WT) *C. elegans* N2, Bristol variety; VC606 *chd-7(gk290) I.* All strains were maintained at 20°C.

To evaluate the percentage of bag-of-worms (Trent *et al*, 1983), synchronized animals were transferred onto an NGM agar plate. Animals with a bag-of-worm phenotype were counted at a Leica dissecting microscope (Leica Mikrosysteme Vertrieb GmbH, Wetzlar, Germany) after 96 h from hatching and removed. The count was repeated after 120 h and 144 h and the total number of altered animals aggregated. To perform the evaluation of the egg-laying (Schafer, 2005), synchronized L4 larval stage animals of each strain were transferred onto NGM agar plates containing a thin bacterial lawn, allowed to lay eggs and transferred to a new plate every 24 h, for 4 days. The eggs laid were counted every day and presented as average of eggs laid every 24 h by a single animal or aggregated as total number of eggs laid by a single animal in 96 h.

#### Drug screening in *C. elegans*

For drug testing, 10 µl of each compound, 10 mM in DMSO 100% from the Chromatin Modification Compound Library (TargetMol, Wellesley Hills, MA, USA; cat. L8300) were dissolved in 133µl of M9 buffer 1× (Stiernagle, 2006) to reach the concentration of 700 µM. 10 µl of every compound concentrated 700 µM was added to each well of a 96-well plate (Corning, Glendale, AZ, USA; cat. 353072), with 24 compounds tested in each 96-well plate. The compounds were immediately read at a Bio-Rad Benchmark Microplate Reader at 595 nm to exclude any significant intrinsic absorbance at this wavelength (Gomez-Amaro *et al*, 2015). *E. coli* strain OP50 for in liquido culturing of *C. elegans* was prepared with a standardized protocol: 6 mL O.N. culture of OP50 bacteria (with an Optical Density of 1.80) were centrifuged at 4,500 ×g for 5 minutes, the bacterial pellet was resuspended in 3 mL of M9 buffer 1×, vortexed for 1 h to ensure complete resuspension of the pellet and, finally, 60µl of antibiotic-antimycotic solution (100×) (Merck, Milan, Italy; cat. A5955-100ML) and 3 µl of cholesterol (5 mg/mL) were added to achieve the final concentrations of 2× and 5 µg/mL, respectively, to obtain the Nematode Liquid Growth Media (NLGM). Egg pellets were collected by bleaching (Porta-de-la-Riva *et al*, 2012) and diluted in NLGM to obtain ∼12 eggs per µl. 5µl of eggs (∼60 eggs) were added to each well containing 10µl of each compound together with 55 µl of NLGM (total final volume of 70µl). Then the plate was transferred in an incubator at 20°C on a swinging base for 4 days allowing animals to grow from eggs to adult in liquido. Each compound was tested at 100 µM final concentration in 1% DMSO, and compared to mock treated animals (1% DMSO). Each plate was prepared avoiding using the more external wells and organized in this way: in two wells there were WT animals treated with mock; in other two wells *chd-7(gk290)* animals treated with mock; in one well only M9 buffer 1× used as blank; and in 48 wells *chd-7(gk290)* animals treated with 24 different compounds (as duplicate). Each plate was replicated three to four times to obtain a final number of five to eight replicates per compound. The absorbance of each well was measured every day, using the Microplate Reader after mixing them for 30 seconds. With this approach we were able to screen up to 96 compounds per week in octuplicate. Some molecules caused a decrease of the turbidity compared to mock, suggesting more animals in the wells, hence a possible rescue of the fertility defect. Other molecules caused an increase of the turbidity compared to mock, suggesting less animals in the wells, hence a possible worsening of the fertility defect. In order to normalize the differences between the absorbance of each compound and the respective mock, we calculated the difference (Δ) in turbidity using two different formulas (one for molecules that cause a decrease and another for molecules that cause an increase in the turbidity):

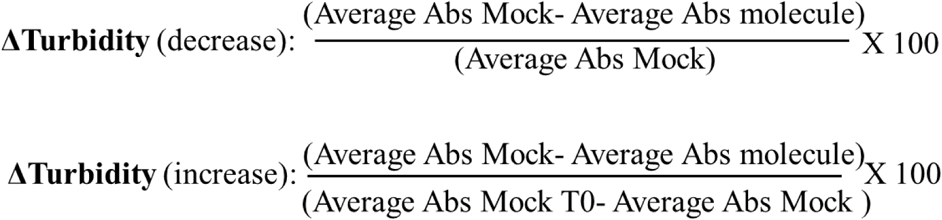

The resulting Δ-turbidity values (for molecules that cause a decrease of the turbidity compared to mock) range from 0 to 100%, where 0 indicates no change in turbidity compared to the mock (no effect) and 100% means, in principle, a molecule that causes a turbidity equal to 0 (no more bacteria) or the most significant rescuing effect. WT animals never reached a 100% decrease in Δ-turbidity, as a complete absence of bacteria in the wells was never observed. For molecules that caused an increase in turbidity compared to mock, the Δ-turbidity values ranged from 0 to negative percentages, with 0 representing a molecule that did not change the turbidity compared to the mock (no effect), and negative values indicating molecules that caused an increase in turbidity (more bacteria) compared to the mock.

### Statistical analysis

In vitro data are presented as mean ± standard deviation (SD). Statistical significance was evaluated using the Prism software (v.9.5.0; GraphPad Software, San Diego, CA, USA). In vivo data are shown as mean ± standard error of the mean (SEM). Statistical analysis has been conducted by using the Prism software for Windows (v.9.0.0; GraphPad Software), except for Figure 4A, where the statistical significance of the difference between two proportions was assessed using a two-sample non-parametric Z-test performed with Microsoft Excel. Number of experiments (N), number of replicates within each experiment (n) and statistical tests used are provided in the corresponding figure legends. All experiments were repeated at least three times. Differences were considered significant whit *p-value* < 0.05.

## Data availability

The data that support the findings of this study are available from the corresponding authors upon reasonable request. RNA sequencing data are available in Gene Expression Omnibus at https://www.ncbi.nlm.nih.gov/geo/, reference number GSE324639 (reviewer token: axcjwsoybxqvbkn).

## Author contributions

**Federica Amoruso**: Data curation; Formal analysis; Validation; Investigation; Methodology; Writing–original draft; Writing–review and editing. **Federica La Rocca:** Data curation; Formal analysis; Validation; Investigation; Methodology; Writing–original draft. **Pamela Santonicola**: Formal analysis; Investigation. **Alyssa J.J. Paganoni**: Formal analysis; Investigation. **Giuseppina Zampi:** Formal analysis. **Stefano Manzini**: Data curation; Visualization. **Fabrizio Fontana**: Methodology. **Riccardo Cristofani**: Methodology. **Roberto Oleari**: Data curation; Methodology; Visualization; Writing–review and editing. **Elia Di Schiavi, Anna Cariboni**: Conceptualization; Resources; Data curation; Formal analysis; Supervision; Funding acquisition; Validation; Investigation; Visualization; Methodology; Writing–original draft; Project administration; Writing–review and editing.

## Ethics Declarations

The authors declare no competing interests.

## Acknowledgements

The N2 wild-type and VC606 *chd-7(gk290) I* nematode strains used in this work were provided by the *Caenorhabditis* Genetics Center, CGC (funded by NIH Office of Research Infrastructure Programs P40 OD010440). The authors thank Wormbase (Sternberg *et al*, 2024). They also would like to acknowledge: Cristina Ruberti and Alessandro Fantin (Università degli Studi di Milano, Milan, Italy) of the Advantage Technology Platform of the Department of Biosciences, for their support in data acquisition and analysis using the ImageXpress Micro Confocal instrument; Paolo Bergamo; Antonio Suppa and Marco Petruzziello (CNR-IBBR, Naples, Italy) for technical support; Chiara Nobile (CNR-IBBR, Naples, Italy) for administrative support. Some of the figures were prepared using <u>Biorender.com</u> (agreement #SQ29JBVMXG).

## Expanded View Figures and Figure Legends

**Figure EV1.**
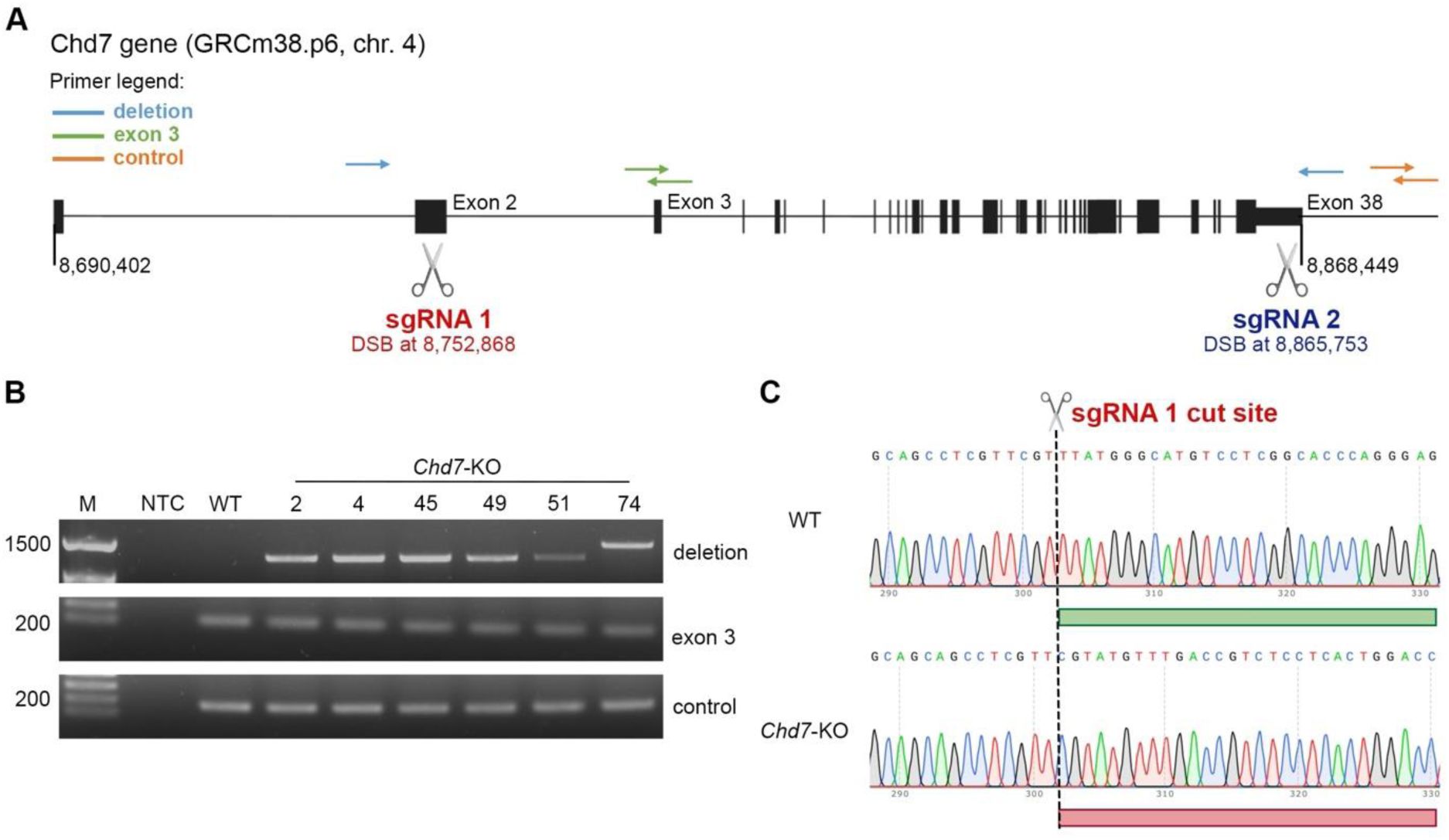
Molecular validation of CRISPR/Cas9-mediated *Chd7* deletion in GN11 cells. (A) Schematic representation of the PCR strategy used to detect *Chd7* gene deletion following CRISPR/Cas9 editing. Primer pairs amplify: an intronic region downstream of the *Chd7* consensus coding sequence (CCDS; orange arrows), exon 3 to assess internal coding sequence integrity (green arrows), a deletion-specific fragment spanning sgRNA cut sites (blue arrows), which is amplified only upon successful genomic excision. In the absence of gene deletion, the large distance between FW_Deletion and REV_Deletion prevents amplification. (B) Endpoint PCR analysis on gDNA from WT and *Chd7*-KO GN11 cell clones. Amplification with deletion-specific primers flanking the sgRNA target sites detects a 1242 bp band in KO clones, consistent with excision of the intervening *Chd7* genomic region. Amplification of exon 3 (101 bp) confirms the presence of residual internal coding sequences in *Chd7*-KO clones, indicative of monoallelic deletion. Control amplification of an unrelated intronic region (167 bp) downstream of the *Chd7* CCDS is shown. (C) Sanger sequencing of PCR products obtained from WT and *Chd7*-KO clones confirming *Chd7* genomic deletion in edited cells. Abbreviations: CCDS, consensus coding sequence; Chr, chromosome; sgRNA, single guide RNA; DSB, double-strand break; FW, forward; REV, reverse.

**Figure EV2.**
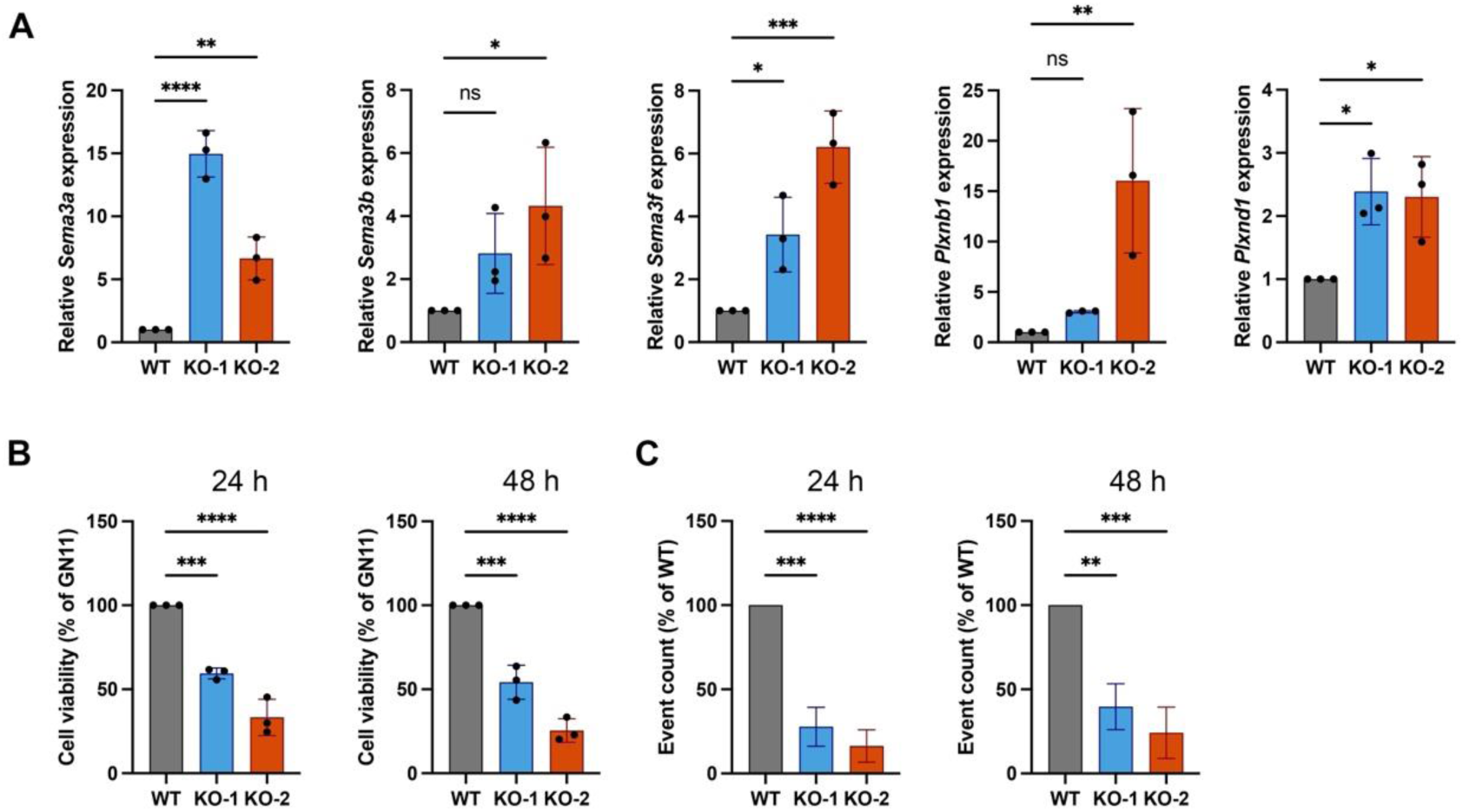
RNAseq validation and cell viability/proliferation analysis in *Chd7*-KO GN11 cells. (A) Expression changes validation by qPCR of indicated *Sema* and *Plxn* transcripts in *Chd7*-KO GN11 cell clones KO-1 (light blue) and KO-2 (orange). Transcript levels were normalized to *Gapdh* and expressed relative to WT controls. Data are shown as mean ± SD (N = 3; One-way ANOVA followed by Dunnett’s multiple comparisons test, * p < 0.05, ** p < 0.01, *** p < 0.001, **** p < 0.0001). (B) MTT-based cell viability assays performed at 24 and 48 h after seeding in WT and *Chd7*-KO GN11 cell clones (KO-1 and KO-2). Data are expressed as % of WT and shown as mean ± SD (N = 3; One-way ANOVA followed by Dunnett’s multiple comparisons test, *** p < 0.001, **** p < 0.0001). (C) Flow cytometry-based quantification of total cellular events at 24 and 48 h after seeding, expressed as % of WT GN11 cells. Data are shown as mean ± SD (N = 3; One-way ANOVA followed by Dunnett’s multiple comparisons test, ** p < 0.01, *** p < 0.001, **** p < 0.0001).

**Figure EV3.**
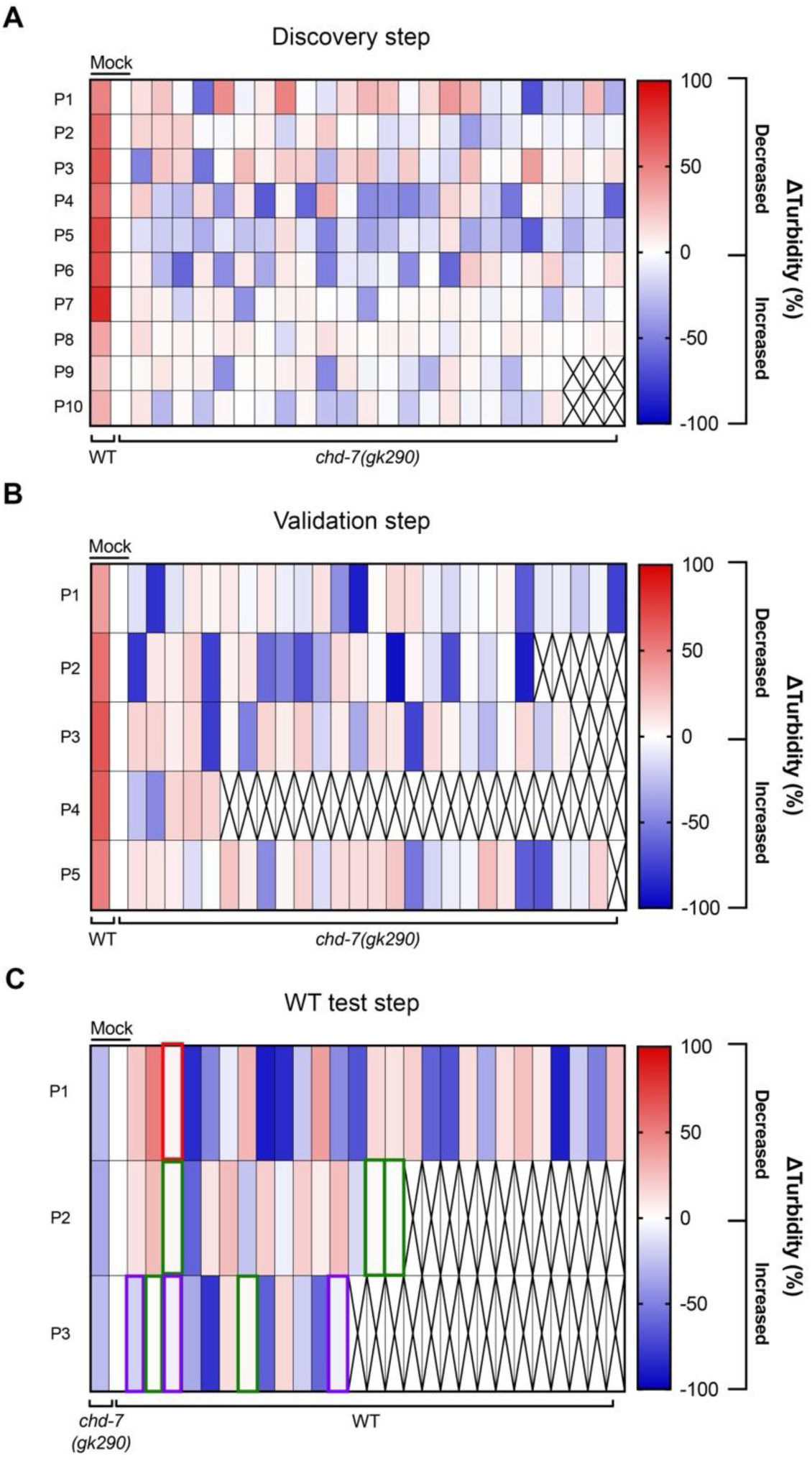
Heatmaps summarizing the results of the discovery (A), validation (B) and WT test (C) screening steps in *C. elegans*. (A) Each horizontal lane represents the data from a single plate read at the Microplate Reader, while each square shows the colour coded average Δ-turbidity of wells filled with individual molecules in the first screening. (B) Heat map summarizing the results of the screening of the best candidates retrieved from the discovery screening. In A and B the first vertical lane displays the results from wells containing WT animals treated with mock (DMSO 1%), while the second vertical lane displays the results from wells containing *chd-7* mutants treated with mock (DMSO 1%); empty cells (with an X) are wells where no molecules have been tested. (C) Heat map summarizing the results of the screening of the best candidates retrieved from the validation screening on WT animals. The first vertical lane displays the results for wells containing *chd-7(gk290)* animals treated with mock (DMSO 1%), while the second lane displays the results for wells containing WT animals treated with mock (DMSO 1%). Green and red squares highlight the molecules that have shown no effect in WT background; purple squares show molecules with an opposite effect in WT compared to *chd-7* mutants. In all heatmaps shades of red represent molecules that decrease turbidity from 0 to 100% (possible rescue), while shades of blue represent molecules that increase turbidity from 0 to -100% (possible worsening).

**Figure EV4.**
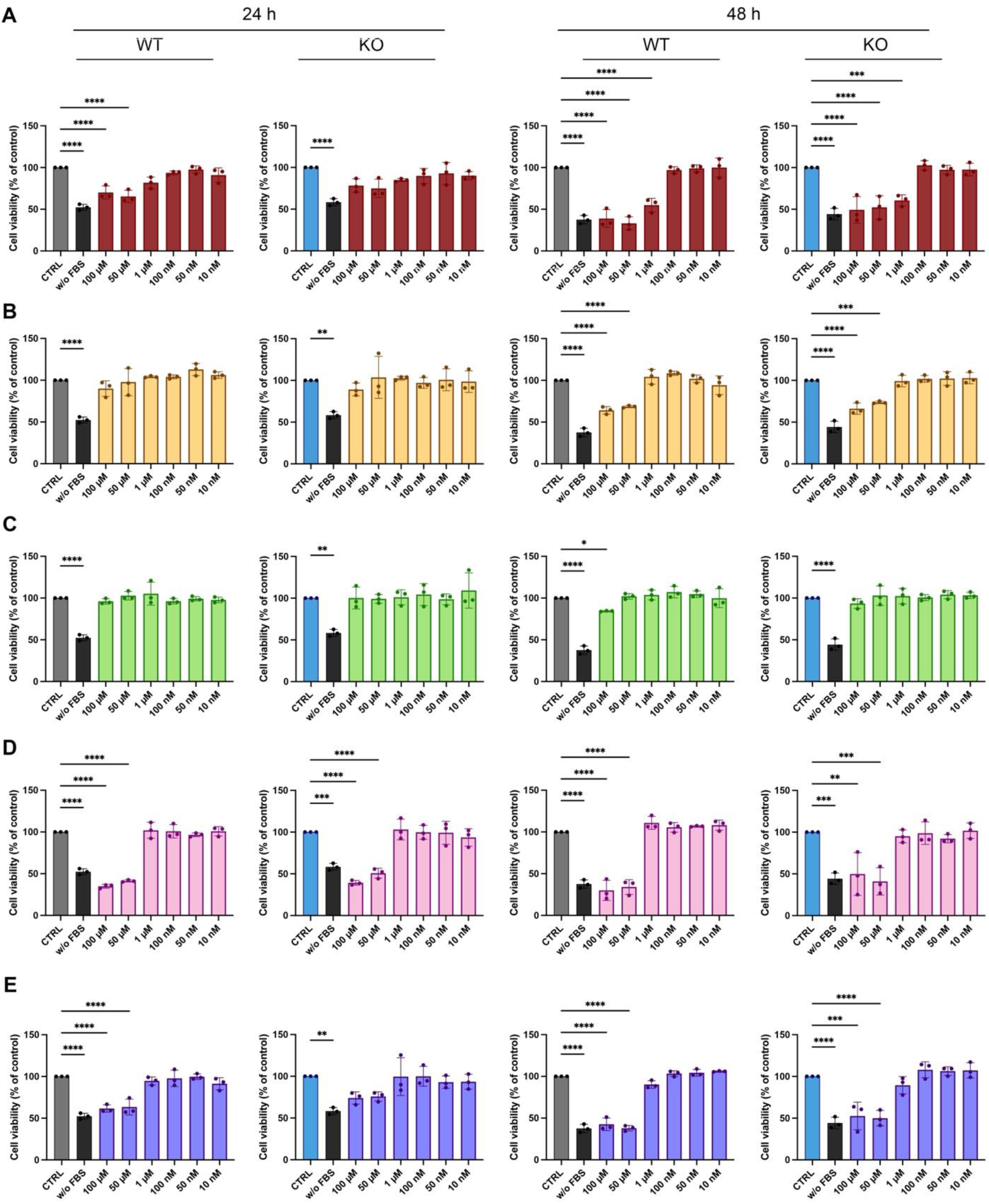
Concentration-dependent cytotoxicity analysis of epigenetic compounds in WT and *Chd7*-KO GN11 cells. (A-E) MTT-based cell viability assays performed after 24 h and 48 h treatment with increasing concentrations (10 nM, 50 nM, 100 nM, 1 µM, 50 µM, 100 µM) of selected epigenetic compounds: (A) Abexinostat in red, (B) BML-210 in yellow, (C) XY1 in green, (D) UNC0646 in pink and (E) HDAC-IN-3 in purple. Each compound was tested in both WT (grey) and *Chd7-*KO (KO-1, light blue) GN11 cells. Cell viability is expressed as a % relative to vehicle-treated controls (CTRL; 0.5% DMSO). Untreated cells maintained in serum-free medium (w/o FBS) were included as negative controls. Data are shown as mean ± SD (N = 3; One-way ANOVA followed by Dunnett’s multiple comparisons test, * p < 0.05, ** p < 0.01, *** p < 0.001, **** p < 0.0001).

**Figure EV5.**
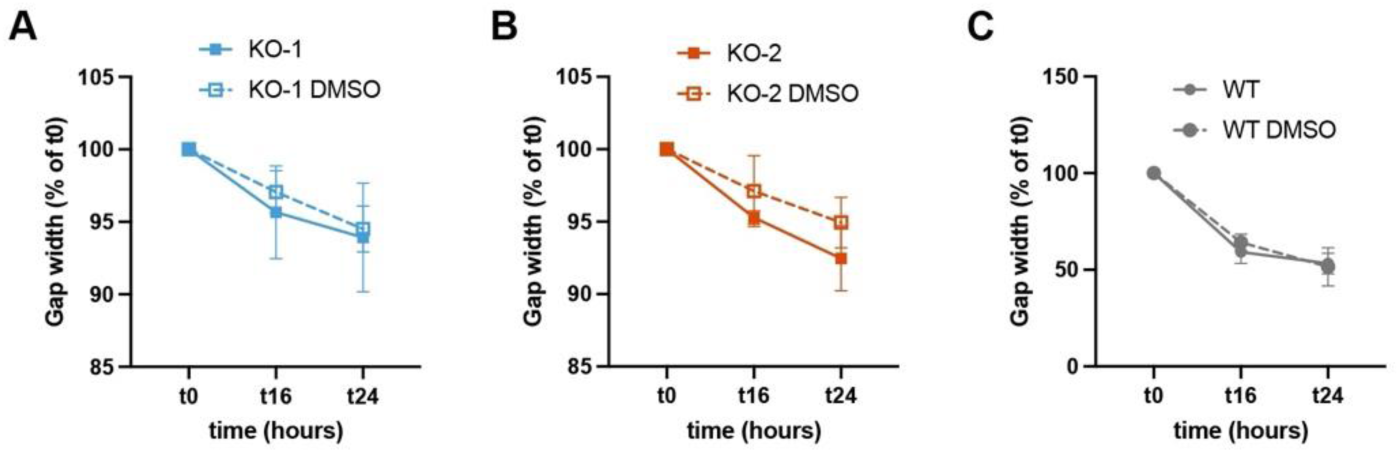
Effect of DMSO exposure on gap-closure dynamics in *Chd7*-KO and WT GN11 cells. (A–C) Quantification of gap width over time (expressed as % of t0) in *Chd7*-KO-1 (A) and KO-2 (B) neuronal cell clones, and WT (C) GN11 cells subjected to wound healing assays. Gap closure was monitored at t0, t16 and t24 and compared to the corresponding DMSO-treated controls (dashed lines). Measurements were normalized to the initial gap width at t0. Data are presented as mean ± SD (N = 3).

